# Role of Rab5 early endosomes in regulating *Drosophila* gut antibacterial response

**DOI:** 10.1101/2022.05.04.490610

**Authors:** Manish Joshi, Annelise Viallat-Lieutaud, Julien Royet

## Abstract

Interactions between prokaryotes and eukaryotes require a dialogue between MAMPs and PRRs. In *Drosophila*, bacterial peptidoglycan is detected by PGRP receptors. While the components of the signaling cascades activated upon PGN/PGRP interactions are well characterized, little is known about the subcellular events that translate these early signaling steps into target gene transcription. Using a *Drosophila* enteric infection model, we show that gut-associated bacteria can induce the formation of intracellular PGRP-LE aggregates which colocalized with the early endosome marker Rab5. Combining microscopic and RNA-seq analysis, we demonstrate that RNAi inactivation of the endocytosis pathway in the *Drosophila* gut affects the expression of essential regulators of the NF-κB response leading not only to a disruption of the immune response locally in the gut but also at the systemic level. This work sheds new light on the involvement of the endocytosis pathway in the control of the gut response to intestinal bacterial infection

**Graphical abstract:** 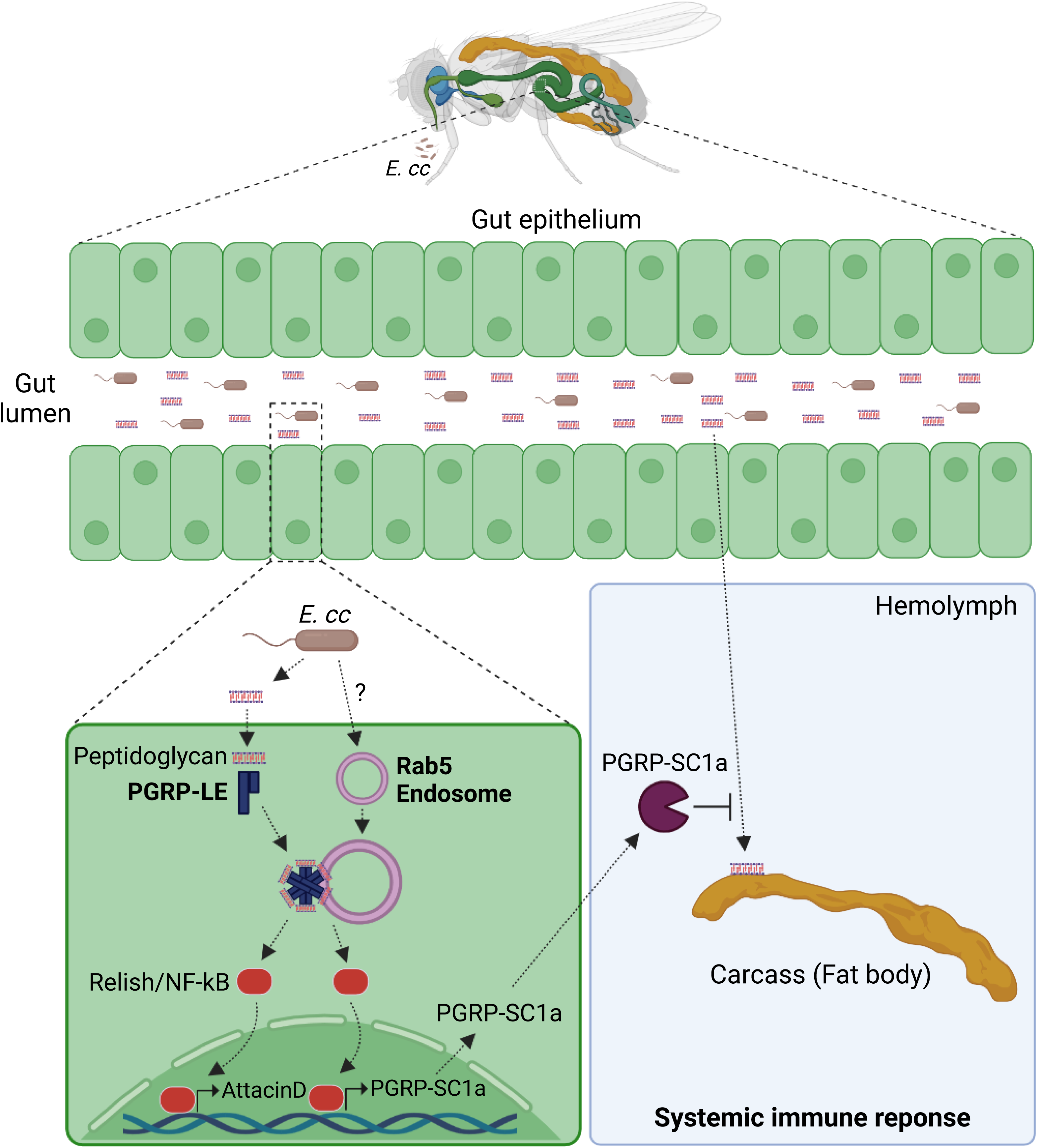

## Introduction

Given that microbes and animals have been evolving together for millions of years, it is not surprising that bacteria profoundly influence host biology, such as the developmental pattern they follow, the type of immunological response they elicit, or even the behavior they exhibit^1^. Some of the interactions between prokaryotes and eukaryotes are mediated by a molecular dialogue between microbe-associated molecular patterns (MAMPs) and host pattern recognition receptors (PRRs)^2^. MAMPs are highly conserved molecular signatures in entire classes of microbes but absent in the host. Their recognition by germline-encoded intracellular and membrane-associated receptors called PRRs triggers *ad hoc* responses in the host^3^. The bacterial peptidoglycan has all the characteristics of a true MAMP^4,5^. Also known as murein, PGN is present in almost all bacteria where it contributes to the shape and integrity of the cell wall. PGN is a glycan polymer composed of alternating residues of β (1,4)-linked N-acetylglucosamine (GlcNAc) and N-acetylmuramic acid (MurNAc) that are cross-linked by short peptide bridges, the exact composition of which is dependent on the bacterial species and growth conditions^4,6^. During bacterial proliferation, up to 50% of the PGN is degraded by a turnover process that generates structurally diverse PGN fragments, called muropeptides^7^. Their recognition by PRRs belonging to various classes of proteins, such as Nod-like receptors (NLRs) and PeptidoGlycan Recognition Proteins (PGRPs) in animals and Lysine-Motif (Lys-M) in plants, is necessary to mobilize host antibacterial defenses, but also to support developmental processes or activate behavior, to name a few^8–13^.

For the past few decades, *Drosophila melanogaster* has served as a model for studying aspects of innate immunity that might otherwise be obscured by the actions of the adaptive immune response^14–17^. Previous work reveals that PGN sensing by PGRP proteins explains many of the interactions that take place between flies and bacteria^18–21^.While Di-amino-pimelic acid PGN (DAP-type PGN) is found in the cell wall of Gram-negative bacteria, Lysine PGN forms the cell wall of Gram-positive bacteria^4^. When bacteria colonize the gut or enter the body cavity of a fly, they release PGN into the gut lumen or into the circulating blood of the insect^22^. Detection of circulating DAP-type PGN by the membrane receptor PGRP-LC expressed on immune cells (adipocytes and hemocytes) triggers NF-κB/Imd-dependent production of immune effectors, such as antimicrobial peptides (AMPs), but also of regulators that control the intensity of the immune response^23–27^. This PGN/PGRP-dependent NF-κB activation also occurs locally in enterocytes that are in direct contact with the gut microbiota. In some cells such as enterocytes, PGN is detected intracellularly by the cytosolic receptor PGRP-LE which shares functional properties with the mammalian PGN sensor Nod2^20,28–32^. Interestingly, genetic data indicate that PGN from the microbiota can reach the insect bloodstream where it activates NF-κB signaling in immunocompetent cells and antagonizes SREBP-dependent lipogenesis^22,33^. More surprisingly, this same metabolite can also reach the brain where it modulates, in an NF-κB dependent manner, the activity of certain neurons, and thus modulate the behavior of infected flies^34–36^.

Mainly through genetic and biochemical approaches, the epistatic relationship between the different components of the NF-κB signaling cascade, activated during PGN detection, has been elucidated in great detail^37^. Via biochemical methods, some of the molecular interactions that take place during Imd signaling pathway activation have been elucidated^37^. It was recently shown that PGRP-LC, PGRP-LE and Imd can form aggregates via their RHIM motifs to promote Imd signaling activity^23^. However, apart from the transcriptional reporters of AMPs^38^, no other molecular tools are available to visualize PGN detection by PGRP and to follow the subsequent activation of downstream NF-κB transduction mechanisms in cells. AMP reporters are only activated in some of the cells that detect PGN and are visible by microscopy much later than the onset of NF-κB activation measured by RT-qPCR on target genes such as AMPs. Furthermore, they are inefficient for tracking PGN-mediated responses that are independent of AMPs. In addition, whereas the components of the immune signaling cascades activated upon MAMPs recognition by PRRs are well known, the subcellular mechanisms by which these proteins transduce the signal remains largely unexplored^39,40^.To further characterize PGN sensing and transduction events, we generated, via CRISPR/Cas9 technology, flies carrying endogenously tagged versions of the cytosolic PGN receptor, PGRP-LE. Microscopic and genetic characterization of these chimeric proteins allowed us to demonstrate the important role played by the Rab5 endosomes in regulating the response of the fly gut to bacterial infection. These results show that, as previously demonstrated for the mammalian PGN sensor NOD2, components of the early endocytic pathway are directly involved in transducing immune signals in flies^39,40^.

## Results

### Generating tools to visualize PGRP-LE-dependent NF-κB signaling *in vivo*

To visualize PGN sensing and signaling events *in vivo*, and at the cellular and subcellular levels, we created transgenic flies in which the PGRP-LE sensor is fused to either a V5 epitope or an eGFP protein (Figure 1A). While the V5 tag was inserted between the PGRP domain and the cRHIM motif of PGRP-LE, the eGFP protein was added to the C-terminus of the PGRP-LE locus (Figure 1A). In these flies, transcription of the PGRP-LE::V5 or PGRP-LE::GFP loci is regulated by the endogenous promoter and all PGRP-LE proteins produced *in vivo* are labeled. To test whether these proteins can be detected *in vivo*, guts of PGRP-LE::V5 and PGRP-LE::GFP were orally infected with *Erwinia carotovora carotovora* (*E.cc*), a strong activator of the NF-κB signaling cascade in the gut and in remote tissues ^41^. Staining corresponding to V5 or eGFP was barely detectable in the intestines of uninfected flies (Figures 1B, 1C and S1A). However, in flies fed for 6h with *E.cc*, PGRP-LE::V5 was detected as aggregates throughout the midgut (except in the R5 domain) with a strong accumulation in R4, a domain previously shown to be almost entirely dependent on PGRP-LE with respect to NF-κB pathway activation (Figures 1B-D and S1A) ^20^. These aggregates were also detected in PGRP-LE::GFP infected guts (Figure S1B). To ensure that V5 staining corresponded only to PGRP-LE protein, staining was performed in cell clones in which PGRP-LE was downregulated by RNAi. In cells in which PGRP-LE mRNA levels were effectively reduced, PGRP-LE aggregates were no longer observed, in contrast to neighboring wild-type cells (Figures 1E and S1C). Furthermore, infected guts in which PGRP-LE was ubiquitously downregulated (using the Mex-Gal4 driver), no longer activated the transcription of *AttacinD,* a well characterized readout of PGRP-LE mediated NF-κB activation in the posterior midgut (Figures S1D and S1E). To test whether the accumulation of aggregates was a consequence of increased PGRP-LE transcription after infection or transgene-related, PGRP-LE mRNA levels were quantified in wild-type, PGRP-LE::V5, and PGRP-LE::GFP strains. In none of these genetic conditions were the PGRP-LE transcript levels different (Figure 1F). This suggests that enteric bacterial infection triggers aggregation of PGRP-LE proteins already present in gut cells before infection. We next tested whether the tags added to PGRP-LEs affect their ability to detect and transduce the bacterial signal. As shown in Figure 1F, the inducibility of *AttacinD* transcripts upon *E.cc* oral infection remains similar in wild-type (w-), PGRP-LE::GFP, and PGRP-L::V5 flies. Using cell-specific markers, we showed that PGRP-LE aggregates are formed in intestinal stem cells, enterocytes, and entero-endocrine cells (Figures S1F-H). Overall, these data demonstrate that the tools we created to visualize PGRP-LE *in vivo* did not alter the PGRP-LE-dependent immune response in the gut and that oral *E.cc* infection triggers PGRP-LE aggregation in gut cells.

**Figure 1:**
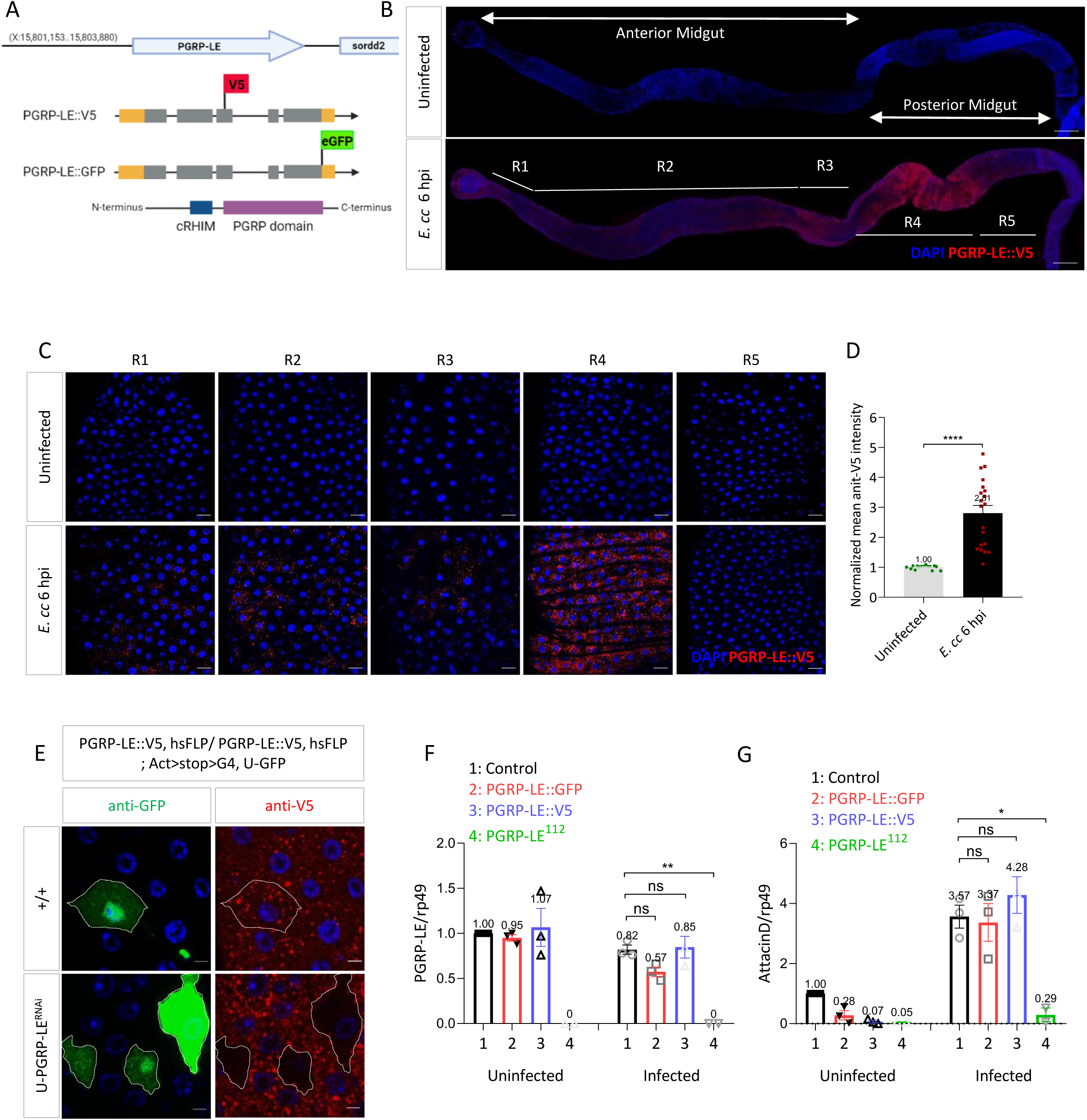
Gut *E. cc* infection induces PGRP-LE aggregation. **(A)** Schematic representation of PGRP-LE locus. The PGRP-LE::V5 and PGRP-LE::GFP were generated by inserting V5 (between PGRP-domain and cRHIM motif) and eGFP (at C-terminus) respectively using CRISPR-Cas9 method. The exonic coding sequences are depicted in grey and non-coding sequences are shown in orange. V5 tag is shown in red and eGFP tag is marked with green. The PGRP domain and cRHIM motif in protein are shown in blue and magenta, respectively. **(B)** Adult midguts from uninfected and infected flies. Immunofluorescence showing PGRP-LE::V5 in red and DNA in blue. Different regions of the midgut marked from R1 to R5. Scale bar represents 200 µm. **(C)** Confocal images of different midgut regions (R1 to R5) showing PGRP-LE::V5 in red and DNA in blue from uninfected and infected flies. Scale bars represent 20 µm. **(D)** Mean fluorescence intensity of PGRP-LE::V5 staining quantified in R4 region from five midguts. Values were normalized with uninfected controls and plotted as mean ± SEM. ****p < 0.0001; Mann-Whitney test. **(E)** Confocal images of control and PGRP-LE^RNAi^ clones from R4 region of posterior midguts from infected flies showing GFP (clones) in green, PGRP-LE::V5 clusters in red and DNA in blue. Scale bars represent 5 µm. (**F** and **G**) mRNA levels of *PGRP-LE* (F) and *AttacinD* (G) of indicated genotypes in posterior midguts of uninfected and infected flies from three independent experiments with ten females per genotype per experiment. ns (non-significant), p > 0.05; *p < 0.05; ***p <0.001; One-way ANOVA with Dunnett’s post-test. Flies were orally infected with *E. cc* for 6 hours. The experiment was repeated three times. Confocal images of one representative experiment are shown. For RT-qPCR results, mRNA levels in uninfected control flies were set to 1, values obtained with other genotypes were expressed as a fold of this value as mean ± SEM.

### PGRP-LE aggregation correlates with activation of the NF-κB pathway

To characterize PGRP-LE aggregates, time course oral infections were performed. As early as 1h after infection, PGRP-LE aggregates were detected in enterocytes of R4 region (Figures 2A and 2B). The size and number of PGRP-LE aggregates increased over time. However, whereas their number peaked at 6hpi and then decreased, they continuously increased to a maximum size at 24hpi (Figures 2A-C). Interestingly, the kinetics of PGRP-LE aggregation paralleled that of the NF-κB signaling pathway quantified by *AttacinD* transcription (Figures 2B and 2D). We then monitored aggregate formation in guts infected with *E. coli* a poor inducer of intestinal NF-κB signaling. Guts orally infected with *E. coli* did not show any sign of PGRP-LE aggregation and of NF-κB pathway (Figures 2E-G). Guts colonized with Gram-positive bacteria or commensal bacteria such as *Lactobacillus plantarum*^42^, *Acetobacter pomorum*^43^ or the entomopathogenic bacterium *Pseudomonas entomophila*^44^ showed no evidence of PGRP-LE aggregation and were unable to activate *AttacinD* transcription (Figures 2H-J). Interestingly, *E.cc* mutant for the *Evf* gene that has been shown to be required for gut AMP induction, did not induce PGRP-LE aggregation^45^. Collectively, these data demonstrated a strong correlation between the ability of a bacterial species to activate the NF-κB pathway and trigger PGRP-LE clustering.

**Figure 2:**
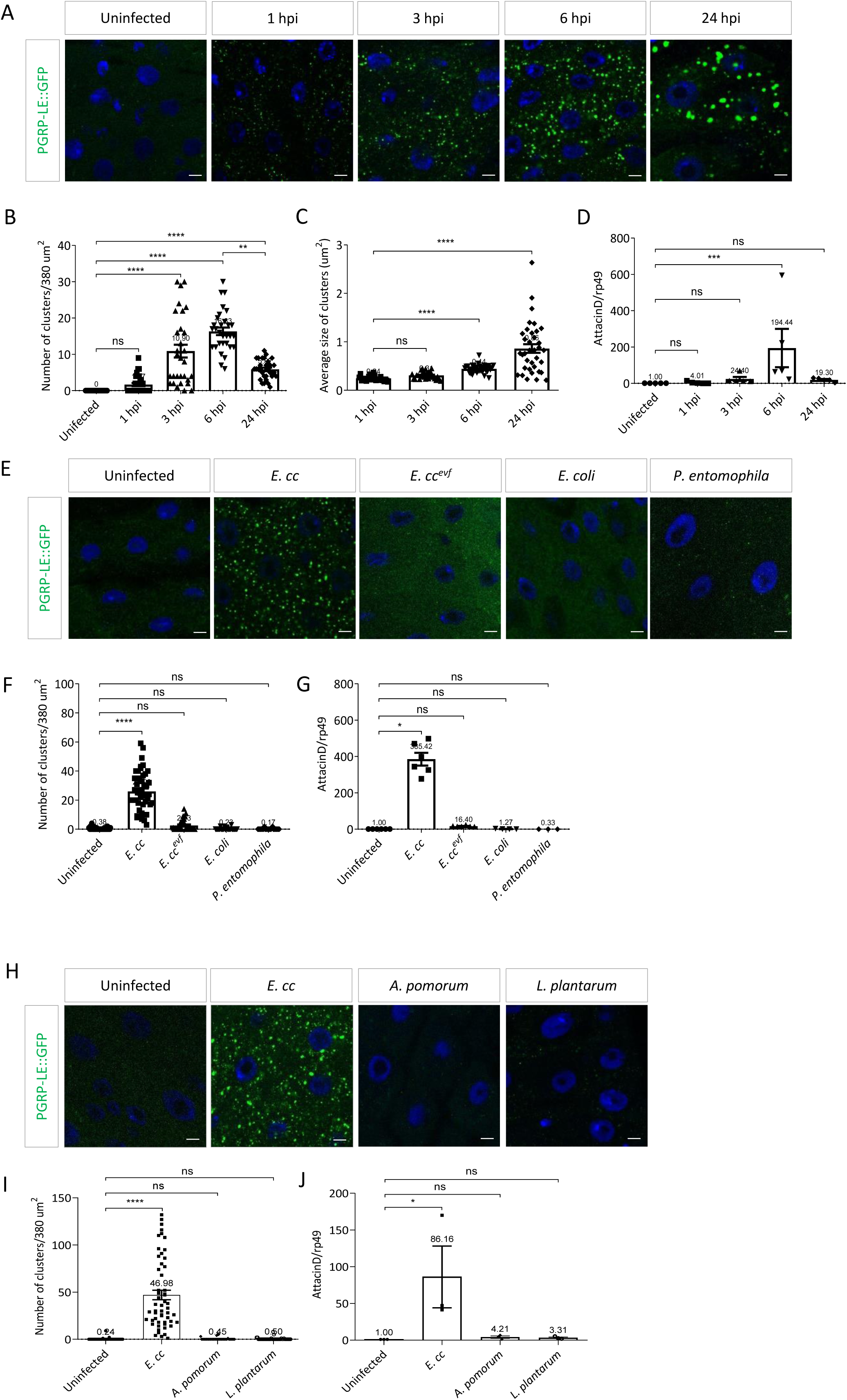
PGRP-LE aggregation correlates with *AttacinD* transcription. **(A)** Confocal images of PGRP-LE::GFP (Green) clusters from R4 region of posterior midgut of uninfected flies and *E.cc* infected flies at different time points. GFP is shown in green and DNA in blue. (**B** and **C**) Quantification of numbers (B) and average size (C) of PGRP-LE::GFP clusters from uninfected and *E. cc* infected flies at different time points from five posterior midguts. ns (non-significant), p > 0.05; ***p < 0.001; ****p < 0.0001; Kruskal-Wallis test. **(D)** *AttacinD* mRNA levels in posterior midguts of uninfected and *E. cc* infected flies at different time points. Results correspond to five independent experiments with ten females per genotype per experiment. Ns (non-significant), p > 0.05; ***p < 0.001; Kruskal-Wallis test. **(E)** Confocal images of PGRP-LE::GFP (Green) clusters from uninfected and infected with *E. cc., E. cc^evf^,E. coli, or P. entomophila*. GFP is shown in green and DNA in blue. **(F)** Quantification of numbers of PGRP-LE::GFP clusters from uninfected flies and flies infected with *E. cc., E. cc^evf^, E. coli, or P. entomophila* from 6 posterior midguts. ns (non-significant), p > 0.05; ****p < 0.0001; Kruskal-Wallis test. **(G)** *AttacinD* mRNA levels in posterior midguts of uninfected flies and flies infected with *E. cc., E. cc^evf^, E. coli, or P. entomophila.* Data correspond to six independent experiments with ten female flies per genotype per experiment. ns (non-significant), p > 0.05; *p < 0.05; Kruskal-Wallis test. **(H)** Confocal images of PGRP-LE::GFP (Green) clusters from R4 region of posterior midguts of uninfected flies and flies infected with *E. cc., A. pomorum or L. plantarum* showing GFP in green and DNA in blue. **(I)** Quantification of numbers of PGRP-LE::GFP clusters from uninfected flies and flies infected with *E. cc., A. pomorum, or L. plantarum* from eight posterior midguts. ns (non-significant), p > 0.05; ****p < 0.0001; Kruskal-Wallis test. **(J)** *AttacinD* mRNA levels in posterior midguts of uninfected flies and infected with *E. cc., A. pomorum or L. plantarum.* Data correspond to three independent experiments with ten female flies per genotype per experiment. ns (non-significant), p > 0.05; *p < 0.05; Kruskal-Wallis test. Flies were orally infected with indicated bacterial species for 6 hours (E-I). The experiment was repeated three times. Confocal images of one representative experiment are shown from R4 region of posterior midguts. Scale bars represent 5 µm. For RT-qPCR results, mRNA levels in uninfected control flies were set to 1, values obtained with other conditions were expressed as a fold of this value as mean ± SEM.

### PGRP-LE^Δ231^ mutant flies do not form aggregates upon *E.cc* infection

Previous work has shown that PGN-derived muropeptides can induce infinite head-to-tail dimers of PGRP-LE and identified key residues required for this oligomerization^23,46^. To test whether such residues are also important in triggering in vivo PGRP-LE aggregation, we generated a PGRP-LE mutant (PGRP-LE^Δ231^::GFP), lacking Glu^231^, a key AA mediating this interaction. Infected guts of such mutant flies no longer showed sign of PGRP-LE aggregation in contrast to controls and were unable to fully activate the Imd pathway (Figures 3A-D). These results suggest that PGN mediated PGRP-LE oligomerization is required to form PGRP-LE aggregates and activate NF-κB pathway.

**Figure 3:**
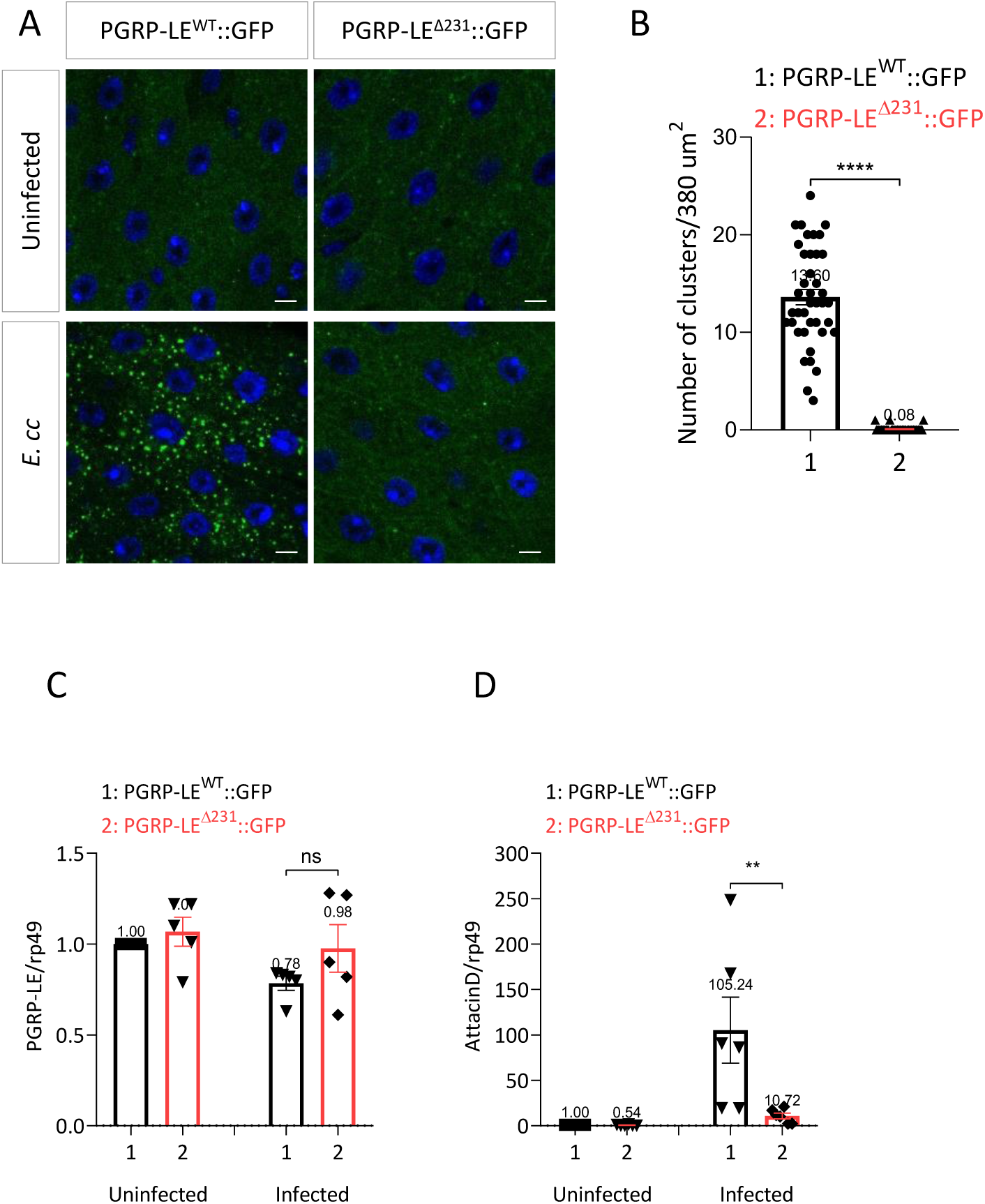
PGRP-LE^Δ231^::GFP mutant do not form aggregates upon *E. cc* infection. **(A)** Confocal images of PGRP-LE::GFP (Green) clusters from R4 region of posterior midgut of PGRP-LE^WT^::GFP and PGRP-LE^Δ231^::GFP from uninfected and infected flies with *E.cc.* PGRP-LE::GFP clusters are shown in Green and DNA is in blue. **(B)** Quantification of numbers of PGRP-LE::GFP clusters from PGRP-LE^WT^::GFP and PGRP-LE^Δ231^::GFP infected flies with *E.cc* from eight posterior midguts. ****p < 0.0001; Mann-Whitney test. (**C** and **D**) mRNA levels of *PGRP-LE* (C) and *AttacinD* (D) of indicated genotypes in posterior midguts of uninfected and infected flies from four independent experiments with ten females per genotype per experiment. ns (non-significant), p > 0.05; **p < 0.01; Mann-Whitney test. Flies were orally infected with *E. cc* for 6 hours. The experiment was repeated three times. Confocal images of one representative experiment are shown from R4 region of posterior midguts. Scale bars represent 5 µm. For RT-qPCR results, mRNA levels in uninfected control flies were set to 1, values obtained with other genotypes were expressed as a fold of this value as mean ± SEM.

### PGRP-LE aggregates colocalize with Rab5

To determine the nature of the PGRP-LE aggregates, we tested their relationship to the subcellular compartments that form the endocytic pathway. Indeed, previous work has show that mammalian NOD1 and kinase RIP2 interact with bacterial peptidoglycan on early endosomes to inflammatory signaling ^47^To this end, *E.cc*-infected guts were stained with antibodies against Rab5 and Rab7, two small GTPases usually associated with early and late endosomes respectively^48,49^. One hour after infection, PGRP-LE aggregates did not colocalize with Rab5 (Figure 4A). Furthermore, the number of detectable Rab5-positive structures increased with time, and at 3hpi most PGRP-LE aggregates were Rab5-positive, reaching almost complete colocalization at 6hpi (Figures 4A and 4B). Strikingly, no Rab5+/PGRP-LE-vesicles were observed, suggesting that Rab5 and PGRP-LE are functionally linked (Figures 4A and 4B). In addition, none of the PGRP-LE vesicles colocalized with the late endosome marker Rab7 at 6 hpi (Figure 4C) or the lysosome marker Lamp1 at 6hpi as well as 24 hpi (Figures 4D and 4E). We next tested the functional relationship between Rab5 and PGRP-LE by analyzing how reducing the level of one will affect the presence of the other. Enterocytes in which PGRP-LE levels were reduced were still able to form Rab5-positive vesicles (Figure 5A). This was also the case for flies carrying PGRP-LE^Δ231^::GFP transgene (Figure 5B). In contrast, enterocytes from *E.cc*-infected guts in which Rab5 was downregulated by RNAi, had more PGRP-LE endosomes and of larger size than those observed in control enterocytes (Figures 5C-F). Similar results were obtained when other components of the endocytic pathway were inactivated. Indeed, RNAi-mediated inactivation of ESCRT complexes members Hrs and VPS28 gave the same phenotype (Figures S4A-C and S4F-H). In contrast, interfering with the Rab7 and lysozyme-associated protein Lamp1 had no effect on PGRP-LE aggregate formation (Figures S2A-D). These results collectively demonstrate that *E.cc* has the specific ability to trigger PGRP-LE/Rab5 vesicle formation in intestinal cells. In addition, these results also show that interfering with the normal function of the endocytic pathway perturbs the formation of these PGR-LE vesicles in *E.cc* infected guts.

**Figure 4:**
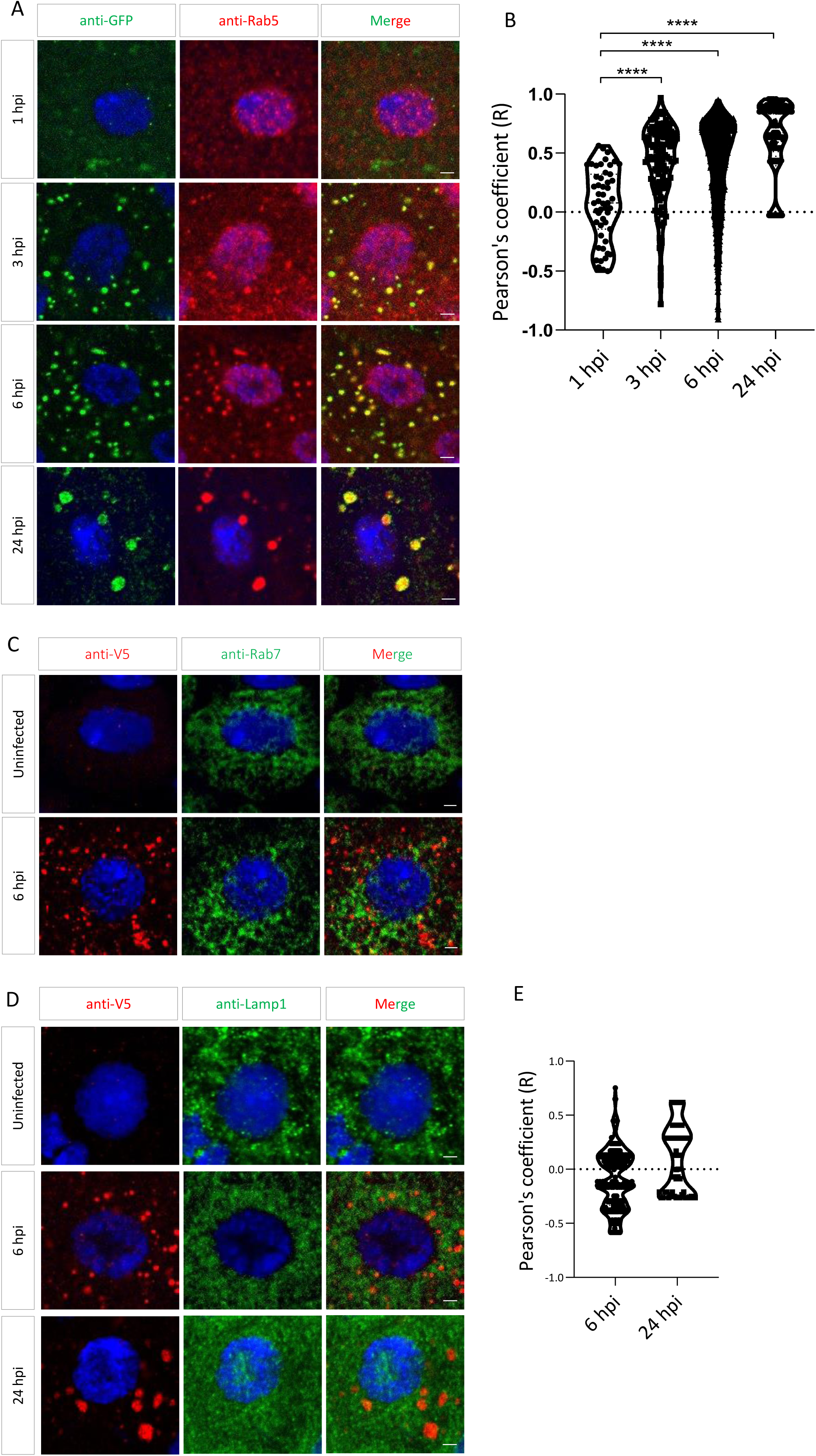
PGRP-LE aggregates are Rab5 positive. **(A)** Confocal images of posterior midguts from *E.cc* infected flies at different time points showing PGRP-LE::GFP clusters in green, Rab5 positive endosomes in red and DNA in blue. **(B)** Quantification of PGRP-LE::GFP clusters colocalization with Rab5 positive endosomes. Pearson’s R coefficient values are plotted in violin plot from eight midguts. ****p < 0.0001; Kruskal-Wallis test. **(C)** Confocal images of uninfected and infected flies showing PGRP-LE::V5 clusters in red and Rab7 positive endosomes in green. DNA is stained with DAPI (blue). **(D)** Confocal images of posterior midguts from *E.cc* infected flies at different time points showing PGRP-LE::V5 clusters in red, LAMP1 positive lysosomes in green and DNA in blue. **(E)** Quantification of PGRP-LE::V5 clusters colocalization with LAMP1 positive lysosomes. Pearson’s R coefficient values are plotted in violin plot. Flies were orally infected with *E. cc*. The experiment was repeated three times. Confocal images of one representative experiment are shown from R4 region of posterior midguts. Scale bars represent 2 µm.

**Figure 5:**
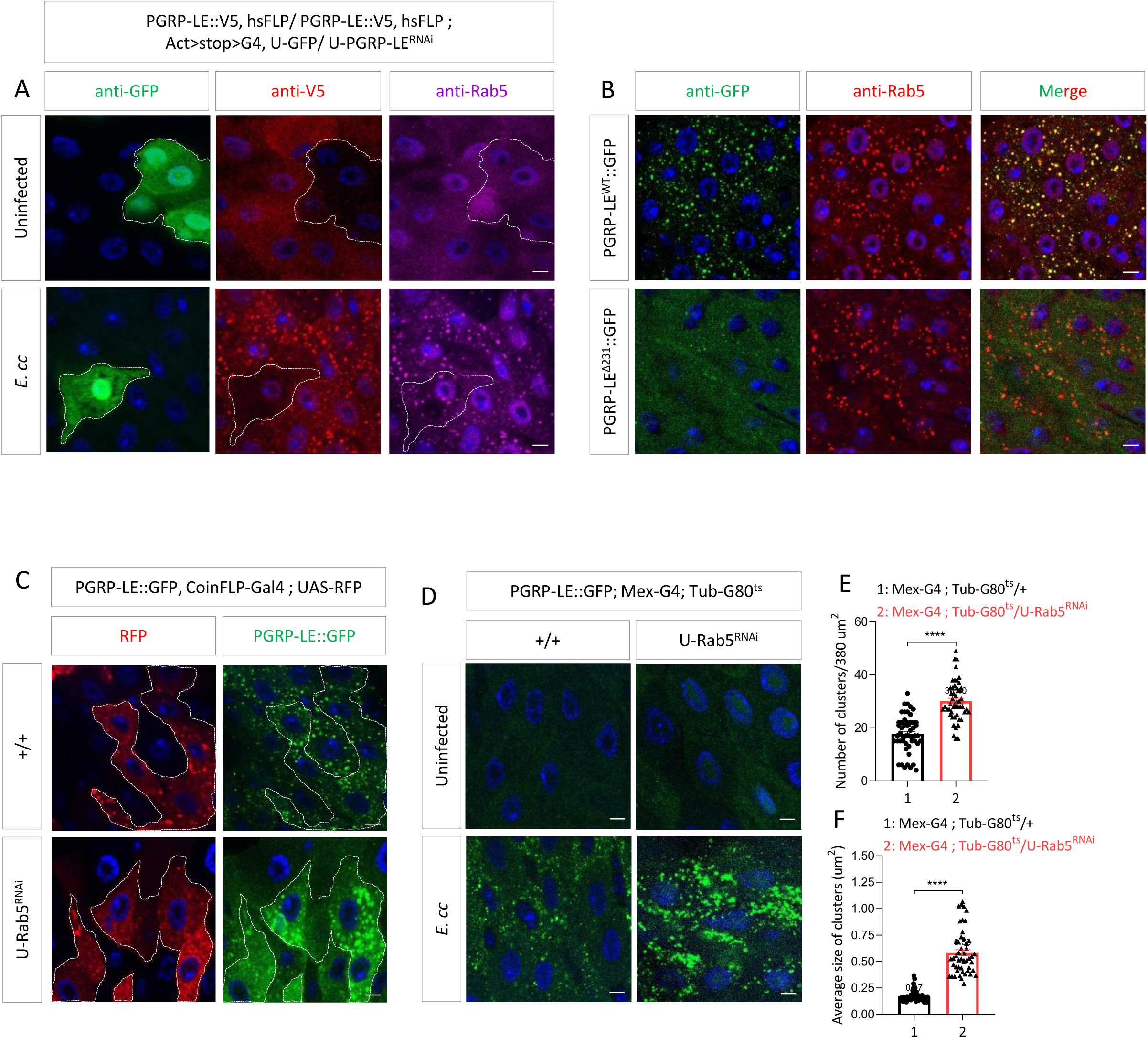
Rab5 endosomes can form in the absence of PGRP-LE. **(A)** Confocal images of PGRP-LE^RNAi^ clones in posterior midguts of uninfected and *E.cc* infected flies. Immunofluorescence showing GFP (clones) in green, PGRP-LE::V5 in red, Rab5 in magenta and DNA in blue. **(B)** Confocal images of PGRP-LE^WT^::GFP and PGRP-LE^Δ231^::GFP in posterior midguts of *E. cc* infected flies showing PGRP-LE::GFP clusters in green, Rab5 positive endosomes in red and DNA in blue. **(C)** Confocal images of control and Rab5^RNAi^ clones in posterior midguts of *E. cc* infected flies showing RFP (clones) in red, PGRP-LE::GFP clusters in green and DNA in blue. **(D)** Confocal images of posterior midguts showing PGRP-LE::GFP clusters in control and Rab5^RNAi^ of infected flies with *E. cc*. GFP is shown in green and DNA in blue. (**E** and **F**) Quantification of numbers (E) and average size (F) of PGRP-LE::GFP clusters in control and Rab5^RNAi^ infected flies with *E. cc* from ten posterior midguts. **p < 0.01; ****p < 0.0001; Mann-Whitney test. Flies were orally infected with *E. cc* for 6 hours. The experiment was repeated three times. Confocal images of one representative experiment are shown from R4 region of posterior midguts. Scale bars represent 5 µm.

### NF-κB and hedgehog signaling components and bacterially derived uracil are not required for the formation of PGRP-LE aggregates

To further characterize the formation of PGRP-LE clusters upon *E.cc* infection, we tested whether components of the Imd pathway were required for the formation of these structures. Cells in which the intracellular transducers Fadd, Dredd, Imd, the PGRP-LC receptor and the Relish transactivator were downregulated by RNAi were still loaded with PGRP-LE aggregates upon *E.cc* infection, suggesting that these components are not required (Figures S2E-G and S3A). Since previous work has demonstrated that *E.cc*-derived uracil modulates DUOX-dependent intestinal immunity via hedgehog-induced Cad99C signaling endosomes, we tested the involvement of bacterial-derived uracil and Hh signaling in PGRP-LE aggregation ^50^. Downregulation of Hh signaling (via inactivation of Smo and Ci) did not interfere with the ability of *E.cc* to induce PGRP-LE aggregate formation (Figure S3B). Similarly, *E.cc* uracil auxotrophic mutants (Ecc^ΔpyrE^), which are unable to induce Cad99C endosome formation, were as effective as wild-type *E.cc* strains at inducing PGRP-LE aggregates (Figure S3C). Furthermore, uracil feeding had no effect on PGRP-LE aggregation (Figure S3C). In addition, DUOX inactivation by RNAi did not interfere with PGRP-LE aggregate formation upon *E.cc* infection (Figure S3E). Therefore, bacterial uracil and DUOX-related gut immunity do not appear to be required for *E.cc*-induced PGRP-LE aggregate formation. We finally tested whether oxidants commonly used to induce stress and tissue damage were sufficient to induce PGRP-LE aggregation ^51^. Although gut cells from paraquat fed flies did not present any sign of PGRP-LE aggregation, they contain Rab5 positive endosomes (Figure S3D). This suggests that two different signals are required to, in one hand, trigger Rab5 endosome formation, and on the other, induce their coating with PGRP-LE proteins.

### Components of early endocytic compartment selectively regulate NF-κB-dependent target genes

Because Rab5 protein associates with PGRP-LE in a kinetic manner parallel to that of NF-κB signaling, we asked whether Rab5 plays a role in PGRP-LE dependent NF-κB signaling. To test this hypothesis, we monitored PGRP-LE-dependent *AttacinD* transcription in the guts of *E.cc*-infected flies in which Rab5 levels were reduced. While inactivation of PGRP-LE by RNAi strongly reduced *AttacinD* mRNA inducibility, this was not the case in guts with reduced Rab5 levels (Figure 6A). Because amidases such as PGRP-LB and PGRP-SC1 are also PGRP-LE-dependent target genes in the gut, we tested their putative regulation by Rab5. In contrast to what was observed with *AttacinD*, PGRP-SC1 transcription was affected not only by reduced PGRP-LE levels but also by Rab5 downregulation in the gut (Figure 6C). This effect was specific since this was not the case for PGRP-LB and PGRP-SC2 (Figures 6D and 6E). Similar results were obtained when Hrs and VPS28 were specifically reduced in gut cells (Figures S4D, S4E, S4I and S4J). Since intestinal amidases have been shown to buffer PGN levels in the intestinal lumen and, consequently, its diffusion into the hemolymph^18,52^, we tested the consequences of Rab5 inactivation in the gut on NF-κB activation in the distant fat body of *E.cc*-infected flies. As previously reported, PGRP-LE activity is required in the gut to prevent NF-κB overactivation in the fat body after oral infection^20^. A similar phenotype was observed in flies with reduced levels of Rab5 in the gut (Figure 6B). To ensure that this phenotype was not secondary to intestinal leakage associated with Rab5 or PGRP-LE inactivation, we performed the “Smurf” test. The “Smurf” assay assesses gut integrity by feeding a blue dye that is impermeable to the intact gut barrier and stays in the gut lumen^53^ Because the blue dye remained confined to the intestinal tract, we concluded that the excessive NF-κB activation in the fat body was not due to intestinal barrier dysfunction (Figures 6F and 6G). Furthermore, whereas downregulation of PGRP-SC1 in the gut did not affect local *AttacinD* transcriptio (Figure 6H), it mimics the increased immune response of the fat body observed upon reduction of PGRP-LE and Rab5 levels in the gut (Figure 6I). To further confirm that the Diptericin ectopic expression observed in gut with reduced Rab5 levels, we performed rescue experiments. Gut-specific overexpression of PGRP-SC1a was sufficient to strongly reduce the fat body Diptericin ectopic expression of Rab5 reduction (Figure 6J). Thus, the association of PGRP-LE and Rab5 in the gut after *E.cc* exposure is necessary for the local induction of PGRP-SC1 that will impact the activation of the immune response at distance in the fat body tissue. However, this Rab5/PGRP-LE association is not necessary for an appropriate local gut antimicrobial response. These results reinforced a model that the endosome pathway is required to activate only a fraction of PGRP-LE-dependent target genes, such as the PGRP-SC1 amidase. By doing so, it prevents systemic activation of the NF-κB pathway by gut-derived PGN.

**Figure 6:**
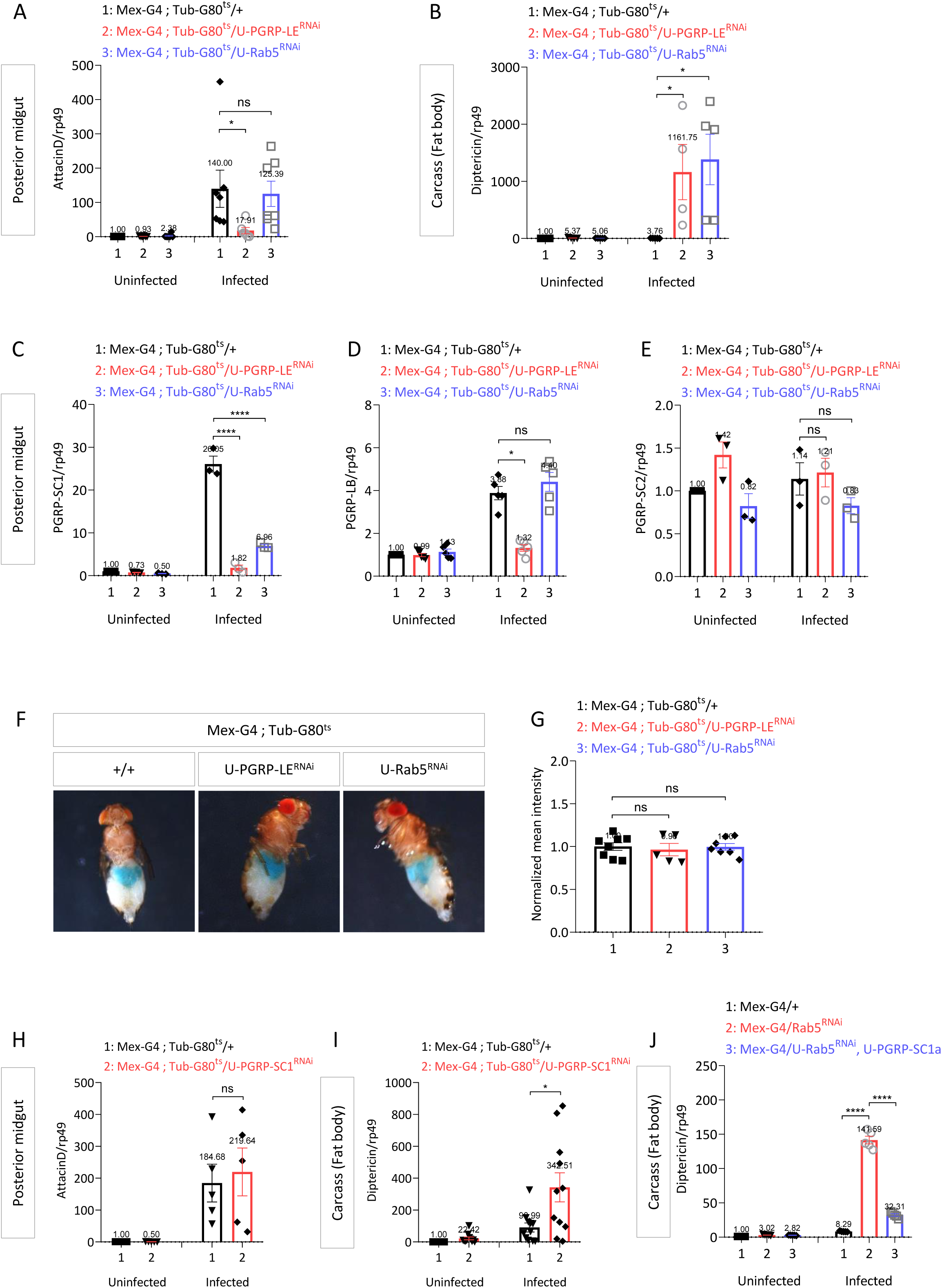
*E. cc*-induced *PGRP-SC1* transcription but not *AttacinD* relies upon Rab5 expression. **(A)** *AttacinD* mRNA levels in posterior midguts of uninfected and *E.cc* infected flies in indicated genotypes. Data correspond to seven independent experiments with ten female flies per genotype per experiment. ns (non-significant), p > 0.05; *p < 0.05; Kruskal-Wallis test. **(B)** *Diptericin* mRNA levels in carcass (fat body) of uninfected and *E.cc* infected flies in indicated genotypes. Data correspond to seven independent experiments with ten female flies per genotype per experiment. *p < 0.05; Kruskal-Wallis test. **(C)** *PGRP-SC1* mRNA levels in posterior midguts of uninfected and *E.cc* infected flies in indicated genotypes. Data correspond to three independent experiments with ten female flies per genotype per experiment. ****p < 0.0001; One-way ANOVA with Dunnett’s post-test. **(D)** *PGRP-LB* mRNA levels in posterior midguts of uninfected and *E.cc* infected flies in indicated genotypes. Data correspond to four independent experiments with ten female flies per genotype per experiment. ns (non-significant), p > 0.05; *p < 0.05; Kruskal-Wallis test. **(E)** *PGRP-SC2* mRNA levels in posterior midguts of uninfected and *E.cc* infected flies in indicated genotypes. Data correspond to three independent experiments with ten female flies per genotype per experiment. ns (non-significant), p > 0.05; Kruskal-Wallis test. **(F)** Intestinal integrity test (Smurf assay) of infected flies with *E. cc* in indicated genotypes. The experiment was repeated three times. Images of one representative experiment are shown. **(G)** Quantification of Smurf assay of infected flies with *E. cc* in indicated genotypes. The experiment was repeated three times. Quantification of Smurf assay of one representative experiment is shown. **(H)** *AttacinD* mRNA levels in posterior midguts of uninfected and *E.cc* infected flies in indicated genotypes. Data correspond to five independent experiments with ten female flies per genotype per experiment. ns (non-significant), p > 0.05; Mann-Whitney test. **(I)** *Diptericin* mRNA levels in carcass (fat body) of uninfected and *E.cc* infected flies in indicated genotypes. Data correspond to 11 independent experiments with ten female flies per genotype per experiment. *p < 0.05; Mann-Whitney test. **(J)** *Diptericin* mRNA levels in carcass (fat body) of uninfected and *E.cc* infected flies in indicated genotypes. Data correspond to five experiments with ten female flies per genotype per experiment. Flies were orally infected with *E. cc* for 6 hours. For RT-qPCR results, mRNA levels in uninfected control flies were set to 1, values obtained with other genotype were expressed as a fold of this value. RT-qPCR results are shown as mean ± SEM.

### Pan-genome effects of Rab5 or PGRP-LE inactivation in posterior midgut following infection

To determine the number and nature of genes whose transcription is dependent on the endocytosis pathway, we used an RNA-seq approach. We compared genome-wide transcript levels in the posterior midguts of wild-type flies and flies in which PGRP-LE or Rab5 were specifically downregulated in the gut. Inactivation of Rab5, but not PGRP-LE, induced downregulation of a cluster (cluster 2) of genes encoding proteins involved in ribosomal function and oxidative phosphorylation (Figures 7A and 7B). As expected, and probably to compensate for the functional inactivation of Rab5, several genes associated with endocytosis processes were specifically upregulated upon Rab5 downregulation (cluster 3) (Figures 7A and 7B). This effect was exacerbated after *E.cc* infection. Since Rab5 and PGRP-LE inactivation impact *AttacinD* transcription differently, we then focused on immune genes regulation by PGRP-LE and Rab5 (cluster 6). We could distinguish three categories. The first includes genes whose inducibility is controlled by both PGRP-LE and Rab5. Confirming the RT-qPCR results above, this group of genes includes *PGRP-SC1* and also another negative regulator of the Imd pathway, PGRP-LF (Figure 7C). Also, as expected, *AttacinD* was found, along with *Pirk,* in a class of genes whose transcription depends solely on PGRP-LE (Figure 7C). Surprisingly, most other immune genes, including all other AMPs except *AttacinD*, were Rab5-dependent but PGRP-LE-independent (Figure 7D). Not only were they still induced upon infection in PGRP-LE^-RNAi^ guts, but their transcriptional inducibility was even higher in PGRP-LE-free guts than in controls. These results showing that most immune genes are regulated independently of PGRP-LE suggest that they are likely regulated by the other upstream Imd pathway receptor, PGRP-LC. Consistently, *PGRP-LC* mRNA levels were lower in Rab5 RNAi samples than in controls (Figure 7D). These data show that by controlling the expression of some essential positive (PGRP-LC) and negative (PGRP-SC1) regulators, the endocytic pathway plays a key role in regulating the gut response to certain bacterial infections.

**Figure 7:**
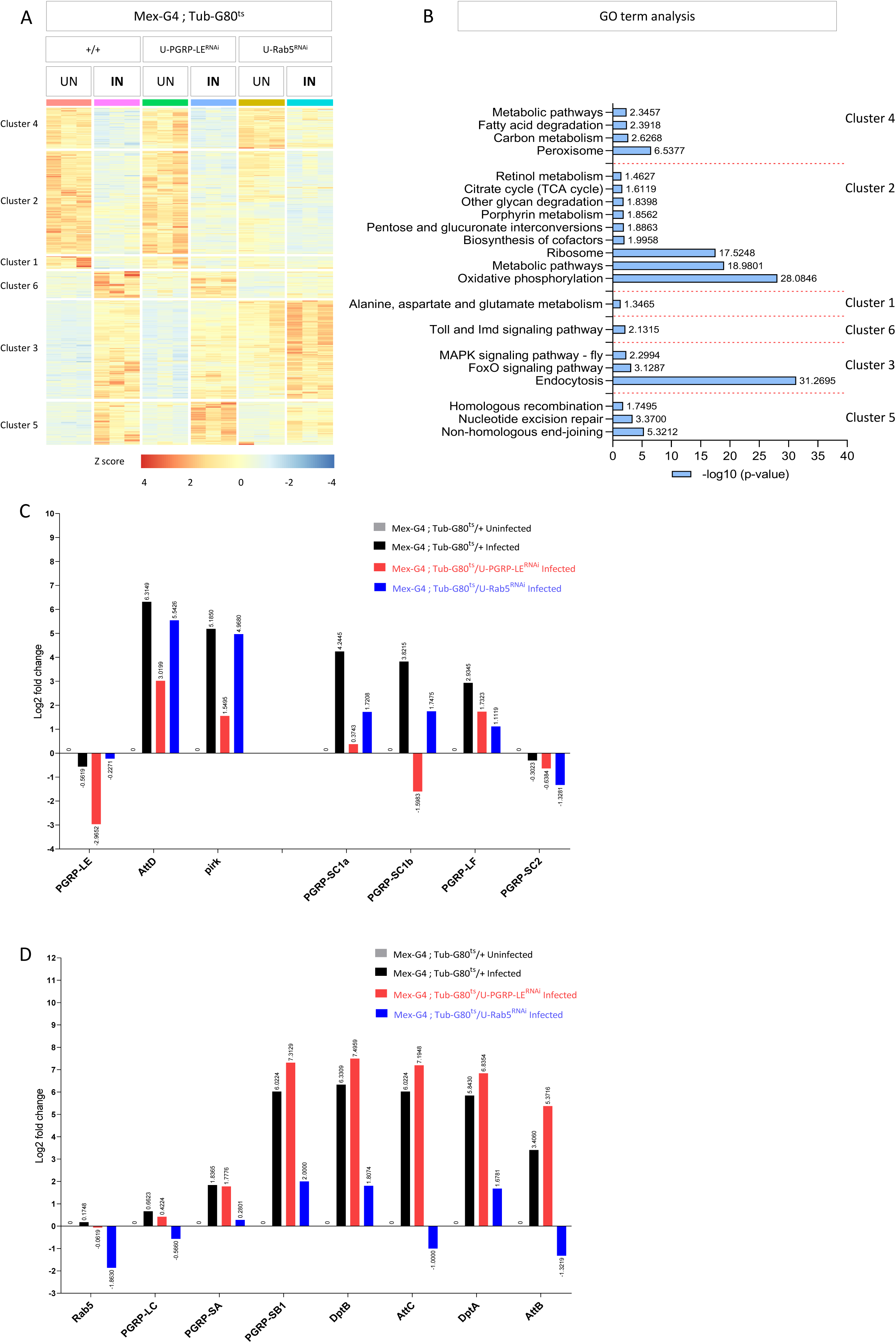
Identification of Rab5 and/or PGRP-LE dependent genes in *E. cc*-infected gut. **(A)** Clustering analysis of the expression of genes from control, PGRP-LE^RNAi^ and Rab5^RNAi^ of uninfected and infected flies. **(B)** Gene ontology (GO) enrichment analysis of different clusters from Fig. 7A (*padj* <0.05). KEGG pathways are shown from GO term. **(C** and **D)** Selected differentially expressed genes (DEGs) of interest from Imd/NF-kB pathway is shown as fold change (*padj* <0.05). For fold change, levels of uninfected control flies were set to 1, values obtained with other genotypes were expressed as a fold of this value. Log2 values are plotted on bar graph.

## Discussion

Using locus-directed tagged-versions of PGRP-LE, we showed here that bacterial gut derived PGN can induce PGRP-LE aggregation in enterocytes. These results are in good agreement with previous data showing that certain muropeptides can induce infinite head-to-tail dimerization of PGRP-LE^46^. Consistently, and in good agreement with the model in which PGN is required to induce PGRP-LE multimerization, flies carrying the PGRP-LE^Δ231^::GFP mutation lose their ability to trigger PGRP-LE aggregation upon *E.cc* oral infection. Interestingly, among all Gram-negative bacteria tested, and thus all having a DAP-type PGN in their cell wall, only *E.cc* was able to induce the formation of PGRP-LE+/Rab5+ vesicles. Furthermore, Rab5 vesicles can be formed in the cells in which PGRP-LE was downregulated, or in cells carrying the PGRP-LE^Δ231^ mutated version of PGRP-LE. Altogether these data suggest a model in which two different *E.cc*-derived signals induce PGRP-LE aggregates, on the one hand, and Rab endosomes, on the other. Consistent with that model, gut exposure to paraquat in the absence of *E.cc*, was sufficient to trigger the formation of Rab5 endosomes that were negative for PGRP-LE. These data demonstrate that, among the bacterial species tested, *E.cc* has the unique property of inducing Rab5/PGRP-LE positive endosomes. The nature of the *E.cc* produced factor triggering Rab5 endosome formation remains to be identified as well as the mechanisms of PGRP-LE recruitment on these endosomes.

Whether PGRP-LE aggregates are related to the previously described amyloid aggregates as components of Imd signaling is unclear and not easily tested^23^. Our attempt to use in vivo dyes to specifically label amyloid was inconclusive. Our data demonstrate that two PGRP-LE-dependent transduction mechanisms coexist in enterocytes. PGRP-LE can signal independently of Rab5 to activate the transcription of some target genes such as *AttacinD* or *PGRP-LB* but requires Rab5 to regulate the production of some target genes such as *PGRP-SC1*. Interestingly, a recent report demonstrated the critical and specific role played by the transcription factor Caudal in regulating the expression of PGRP-SC1 in posterior midgut^54^. Our data suggests that Rab5-containing vesicles may serve as signaling platforms for PGRP-LE-dependent signaling. Interestingly, endosomes have been shown to be specialized platforms for bacterial sensing and Nod2 signaling while the endocytic pathway is required for the other *Drosophila* NF-κB innate immune signaling, Toll^40,50,55^. Other organelles such as mitochondria or peroxisomes have been shown to be signaling platforms^56,57^. In addition, it has been shown that by interfering with Rab11 vesicle exocytosis, the *Drosophila* pkaap protein regulates the secretion of the antimicrobial peptide Drosomycin at the plasma membrane^58^. In addition, recent work has demonstrated that endosome formation is implicated in intestinal epithelial maintenance and ISC-dependent gut renewal^59–61^

Why should such a binary system be required to respond to a single ligand-receptor interaction? It is conceivable that such a system would allow flies to respond to different thresholds of peptidoglycan and thus, potentially, bacterial load. Low levels of PGN would induce AttacinD production via PGRP-LE but independently of Rab5, perhaps via freely diffusible PGRP-LE molecules. Higher levels of PGN, synonymous with high infection, would require the production of a negative regulator to avoid the deleterious and documented effects of NF-κB pathway overactivation already reported^58,62^.

It remains to be understood how the interaction with early endosomes will interfere with downstream signaling. One hypothesis is that some of the components of the Imd pathway are post-translationally modified during their interactions with the endosome. Consistently, *Drosophila* H2Av has been shown to negatively regulate Imd pathway activity by facilitating SUMOylation of Relish^62^.

### Limitations of the study

This study has several limitations. Firstly, it will be useful to test more bacterial species to identify new ones able to trigger PGRP-LE aggregation. This should allow us to identify the mechanisms by which bacteria trigger PGRP-LE aggregation. Secondly, in addition to PGRP-LE, the transmembrane PGRP-LC receptor is playing a key role in regulating enterocytes responses to bacteria. Analyzing PGRP-LC dynamic and localization in regards to the one of PGRP-LE will be very informative to have a broader view on the described phenotype. Finally, it will be important to demonstrate whether or not PGRP-LE aggregates correspond to the amyloid structure previously described.

## Acknowledgments

We thank Emilie Avazeri for technical help, Florent Fioriti for help with RNAseq analyses, Léo Kurz and Olivier Zugasti for comments on the manuscript. This work was supported by ANR BACNEURODRO (ANR-17-CE16-0023-01), ANR PEPTIMET (ANR-18-CE15-0018-02), Equipe Fondation pour la Recherche Médicale (EQU201903007783) and Institut Universitaire de France to J.R.

## Author contributions

Conceptualization, J.R.; Methodology, M.J., A.L.V. and J.R..; Investigation, M.J., A.L.V.; Formal analysis, M.J. and J.R.; Writing – original draft, M.J. and J.R.; Writing – reviewing & editing, J.R.; Funding acquisition, J.R.; Supervision, J.R.

## STAR★Methods

### KEY RESOURCES TABLE

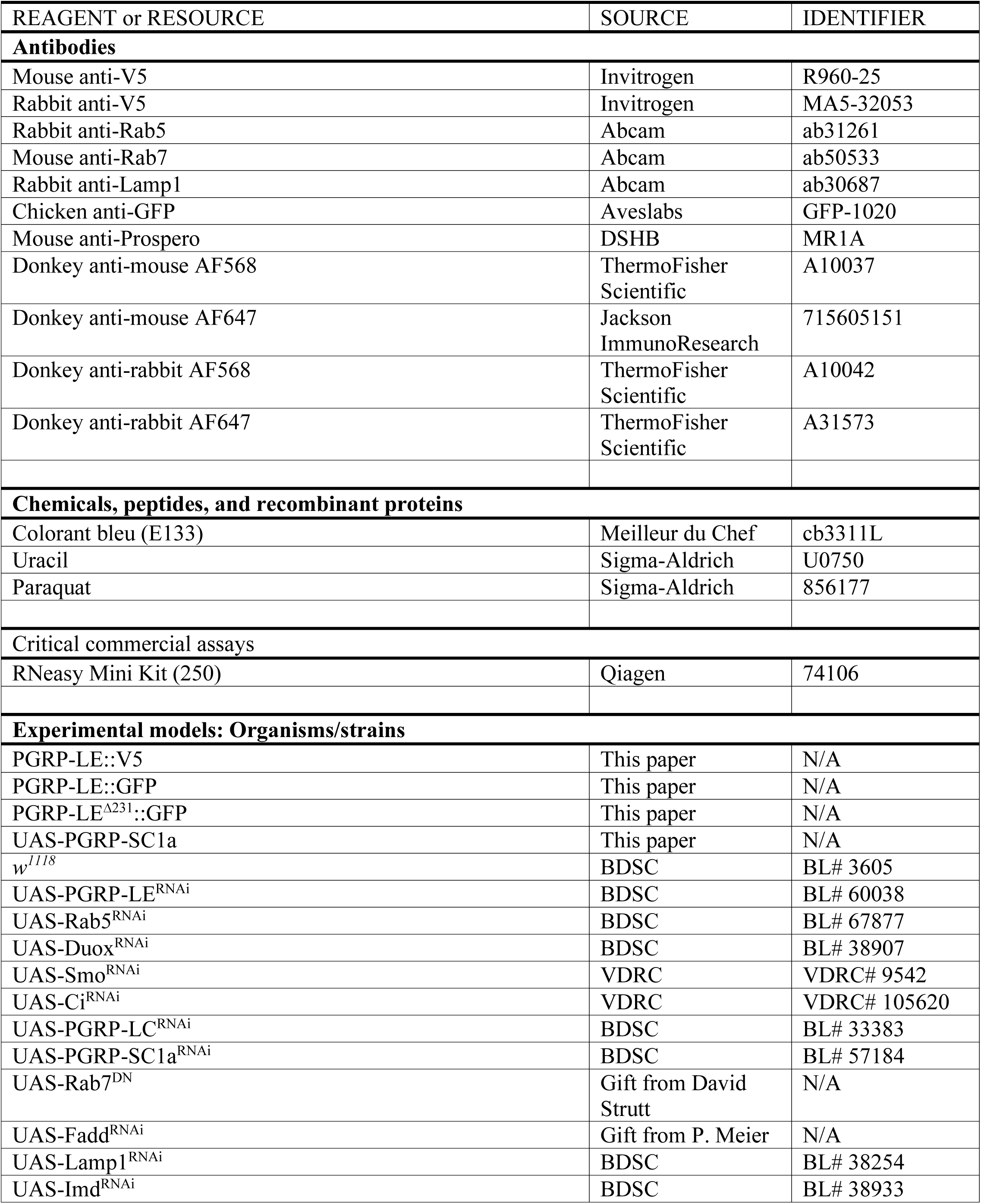
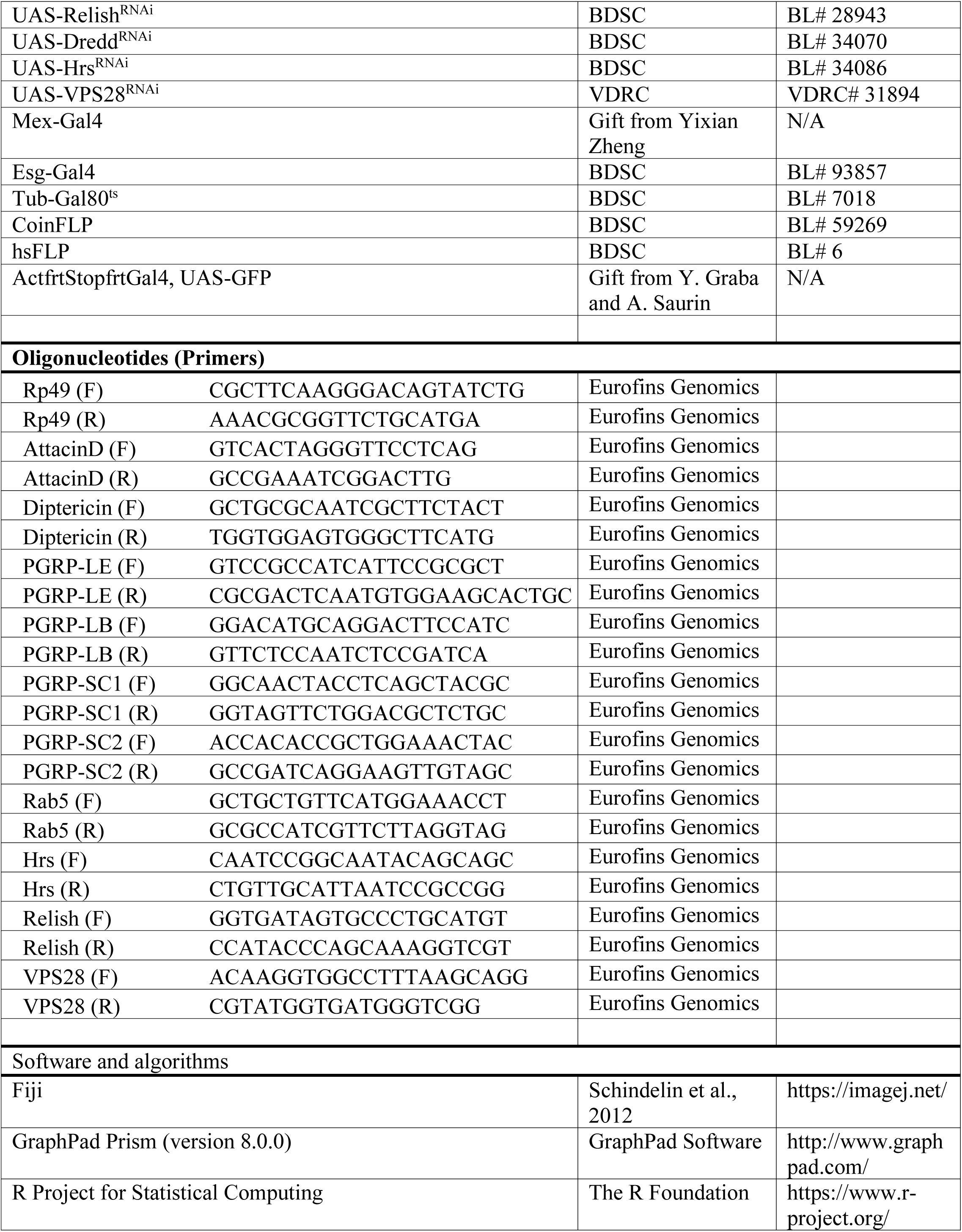

### RESOURCE AVAILABILITY

#### Lead contact

Further information and requests for resources and reagents should be directed to and will be fulfilled by the lead contact, Julien Royet.

#### Materials availability

All fly lines generated in this paper will be made available upon request. A material transfer agreement will be required prior to sharing of materials.

#### Data and code availability

- All data reported in this paper will be shared by the lead contact upon request.
- All codes used to analyze the data in this paper will be shared by the lead contact upon request.
- Any additional information required to reanalyze the data reported in this paper is available from the lead contact upon request.

### EXPERIMENTAL MODEL AND SUBJECT DETAILS

#### *Drosophila* culture media

Adult female *Drosophila melanogaster* was used to perform all the experiments. Flies were grown at 25°C on a yeast/cornmeal medium in 12h/12h light/dark cycle-controlled incubators. For 1 L of food, 8.2g of agar (VWR, cat.#20768.361), 80g of cornmeal flour (Westhove, Farigel maize H1) and 80 g of yeast extract (VWR, cat.#24979.413) were cooked for 10 min in boiling water. 5.2g of Methylparaben sodium salt (MERCK, cat.#106756) and 4 ml of 99% propionic acid (CARLOERBA, cat.#409553) were added when the food had cooled down. For antibiotic treatment (ATB), standard medium was supplemented with Ampicillin, Kanamycin, Tetracyclin and Erythromycin at 50 μg/ml final concentrations.

#### Fly strains and genetics

The following strains were used in this work: PGRP-LE::V5 (This work), PGRP-LE::GFP (This work), PGRP-LE^Δ231^::GFP (This work), UAS-PGRP-SC1a (This work), *w*^1118^ (BL# 3605) Mutant line: PGRP-LE^112^, RNAi-Lines: UAS-PGRP-LE^RNAi^ (BL# 60038), UAS-Rab5^RNAi^ (BL# 67877), UAS-Duox^RNAi^ (BL# 38907), UAS-Smo^RNAi^ (VDRC# 9542), UAS-Ci^RNAi^ (VDRC# 105620), UAS-PGRP-LC^RNAi^ (BL# 33383), UAS-PGRP-SC1a^RNAi^ (BL# 57184), UAS-Rab7^DN^ (Kindly provided by David Strutt), UAS-Fadd^RNAi^ (Kindly provided by P. Meier), UAS-Lamp1^RNAi^ (BL# 38254), UAS-Imd^RNAi^ (BL# 38933), UAS-Dredd^RNAi^ (BL# 34070), UAS-Relish^RNAi^ (BL# 28943), UAS-Hrs^RNAi^ (BL# 34086), UAS-VPS28^RNAi^ (VDRC# 31894). Gal4 Lines: Mex-Gal4 (kindly provided by Yixian Zheng), Esg-Gal4 (93857), Tub-Gal80^ts^ (BL# 7018) Clonal Analysis Lines: CoinFLP (BL# 59269) hsFLP (BL# 6), hsFLP; ActfrtStopfrtGal4, UAS-GFP (Kindly provided by Y. Graba and A. Saurin lab).

Flies of genotypes containing the Gal4/UAS/Gal80^ts^ constructs were reared at 21 °C and transferred to 29 °C for 48 hours before infection to allow the activity of the Gal4 transcription factor.

The RNA*i* stocks were tested for their knockdown efficiencies by using a Mex-Gal4 driver to express these lines, followed by isolation of total mRNA from the posterior midgut, subjecting them to RT-qPCR analysis with respective primers. RNA*i* knockdown efficiencies of the respective lines are as followed: UAS-PGRP-LE^RNAi^ (∼90%), UAS-Rab5^RNAi^ (∼57%), UAS-Hrs^RNAi^ (∼62%), UAS-VPS28^RNAi^ (68%), UAS-PGRP-SC1a^RNAi^ (70%) and UAS-Relish^RNAi^ (∼73%).

#### Bacterial species and Infection

Following microorganism were used: *Erwinia carotovora carotovora 15 2141* (*E. cc*) (grown at 30°C), *E. coli* (grown at 37°C), *Acetobacter pomorum* (grown at 30°C)*, Lactobacillus plantarum* strain WJL (grown at 37°C), *Pseudomonas entomophila (grown at 30*°C*), E. cc evf (grown at 30*°C). Microorganisms were cultured overnight in Luria-Bertani (for *E.* cc, *E. cc evf P. entomophila,* and *E*. *coli*) and MRS medium (for *L*. *plantarum* and *A*. *pomorum*).

##### Oral Infection

We used 3–5 days old adult female raised at 25°C in presence of ATB in the food. Flies were starved for two hours before infection for synchronized feeding. Bacterial cultures were centrifuged at 4,000 RPM for 10 min at room temperature. Cells were serially diluted in water and their concentration was determined by optical density (OD) measurement at 600 nm. The food solution was obtained by mixing a pellet of bacterial culture with a solution of 5% sucrose (50/50) and added to a filter disk (final OD_600_ = 100) that completely covered surface of the fly food. Uracil (100 nM) and paraquat (10mM) were orally administered similar as bacterial infection.

### METHOD DETAILS

#### PGRP-LE::V5

The PGRP-LE::V5 fusion protein transgenic fly line was obtained by inserting, *via* CRISPR mediated recombination, a V5-tag cDNA in the 3rd exon of PGRP-LE, upstream of the PGRP domain“TCTdetector”.The V5 single-stranded oligo DNA nucleotides donor (ssODN) was synthetized by Eurofins MWG (GATGCATACACGTTAACCATAGGATACCGATTTCACTTACAGAATTCAAAATACCCAAGagcggtggtaagcctatccctaaccctctcctcggtctcgattctacgggtggcGAGCTGTgCGCCATCATT CCGCGCTCTTCGTGGCTAGCACAGAAGCCCATGGACGAG). The guide RNA (ATGATGGCGCACAGCTCCTT), was cloned into pCFD3–dU6:3 gRNA (Addgene, 49410). w^1118^; attP2(nos-Cas9) embryos were injected with both V5 ssODN donor (100 ng/µl) and pCFD3-gRNA vector (100 ng/µl). G0 were crossed to the balancer stock Sqh/FM7. Each F1 was crossed again to the balancer stock Sqh/FM7. Then its gDNA was extracted and screened by PCR for V5 presence. Positive lines were confirmed molecularly by sequencing.

#### PGRP-LE::GFP

A PGRP-LE::GFP fusion protein transgenic line was obtained by inserting, *via* CRISPR mediated recombination, the meGFP cDNA at the C-term end of the PGRP-LE protein. The vector donor BS PGRP-LE::GFP was obtained by cloning the meGFP cDNA flanked by 1kb of PGRP-LE homology arms into the Bluescript vector using the following primers:

fw5’arm:CGGGCTGCAGGAATTCATAAGCAACTCCACGAACGT,
rv5’arm:CTCGCCCTTGCTCACTTGTTCCTCCTCCTCGATATTG;
fw3’arm:CAAGCACCGGTCCACGTGAGGGACAAAAGAAGAGCAC,
rv3’arm:GGGCCCCCCCTCGAGTGACCAAACGAATGCAGGAC.

A co-CRISPR strategy (targets: PGRP-LE and ebony e) was used to simplify the screening process^63^. Guide RNAs (PGRP-LE:TCGAGGAGGAGGAACAATGA; ebony: GCCACAATTGTCGATCGTCA) were cloned into pCFD3–dU6: 3 gRNA (Addgene, 49410). w^1118^; attP2{nos-Cas9} embryos were injected with donor (500 ng/µl BS PGRP-LE::GFP) and guide vectors (100 ng/µl pCFD3 PGRP-LE; 100 ng/µl pCFD3 ebony). G0 were crossed to the double balancer stock def w+/FM6; Sb/TM3Ser, e. Each F1 with ebony body color (chrIII: nosCas9* e / TM3Ser, e) was crossed to the balancer stock Sqh/FM7. Then its gDNA was extracted and screened by PCR for GFP presence. Positive lines were confirmed molecularly by sequencing.

#### PGRP-LE^Δ231^::GFP

To obtain PGRP-LE^Δ231^::GFP, co-CRISPR strategy (targets: *PGRP-LE* and ebony *e*) was used to simplify the screening process^63^. Guide RNAs (PGRP-LE: AGTGCTTCCACATTGAGTCG; ebony: CCACAATTGTCGATCGTCA) were cloned into pCFD3–dU6: 3 gRNA (Addgene, 49410).

The y^1^w^1118^ PGRP-LE::GFP;; attP2{nos-Cas9 y+} embryos were microinjected with guide vectors (pCFD3–dU6:3 gRNA; Addgene # 49410; into which the guide RNAs were cloned) at 100 ng/µl pCFD3 PGRP-LE and 100 ng/µl pCFD3 ebony and the corresponding single strand donor oligonucleotide ssODN at 100 ng/µl. G0 flies were crossed to the double balancer stock def w+/FM6; Sb/TM3Ser, e. Each F1 fly with ebony body color (chrIII: nosCas9* e / TM3Ser,e) was crossed to the balancer stock FM7/ Sqh or Y. Then its gDNA was extracted. PCR and sequencing screenings were performed on each F1 ebony fly in order to identify the potential presence of a deletion of amino acid 231.

#### UAS-PGRP-SC1a

The UAS PGRP-SC1a fly line was obtained by integrating a pUASt-PGRP-SC1a-attB vector in the attP2 site of the y^−^w^−^ *nos*PhiC31 ^y+^; attP2^y+^ fly line (modified from BDSC lines #25709 and #25710), *via* transgenesis.

The UAS PGRP-SC1a construct was generated by amplifying the SC1a ORF by PCR from fly genomic DNA (primers F: AACAGATCTGCGGCCGCATGGTTTCCAAAGTGGCTCTCC, R: AAAGATCCTCTAGAGGTACCCTAGCCAGACCAGTGGGAC) and cloning it by

InFusion® (Takara Bio) into the vector pUASt-attB which has attB docking site, Gal4 binding sites, w^+^ marker (DGRC Stock 1419). The pUASt-PGRP-SC1a-attB vector was injected in the embryos of the y^−^w^−^ *nos*PhiC31 ^y+^; attP2^y+^ fly line. Integration in the attP2 docking site was screened in F1 by the presence of the w^+^ marker. Positive lines were confirmed molecularly by sequencing.

#### RNA extraction, RT-qPCR and RNA-seq

RNA from dissected posterior midgut and abdominal carcass (n=10) was extracted with RNeasy Mini Kit (250). Three hundred nanograms of total-RNA was then reverse transcribed in 20 ul reaction volume using the Superscript III enzyme (Invitrogen) and random hexamer primers. Quantitative real-time PCR was performed on a CFX96 Real-Time PCR Detection System (BIO-RAD) in 96-well plates using the FastStart Universal SYBR Green Master (Sigma-Aldrich). The amount of mRNA detected was normalized to control rp49 mRNA values. Normalized data was used to quantify the relative levels of a given mRNA according to cycling threshold (Ct) analysis. mRNA levels in uninfected control flies were set to 1, values obtained with other genotype were expressed as a fold of this value. RT-qPCR results are shown as mean ± SEM from posterior midgut of ten female flies per genotype from at least three independent experiments.

For RNA-seq analysis, RNA was extracted from dissected posterior midguts with RNeasy Mini Kit (250). Library preparation was performed at the GenomEast platform at the Institute of Genetics and Molecular and Cellular Biology using Illumina Stranded mRNA Prep Ligation - Reference Guide - PN 1000000124518. RNA-Seq libraries were generated according to manufacturer’s instructions from 300 ng of total RNA using the Illumina Stranded mRNA Prep, Ligation kit and IDT for Illumina RNA UD Indexes Ligation (Illumina, San Diego, USA). Briefly, Oligo(dT) magnetic beads were used to purify and capture the mRNA molecules containing polyA tails. The purified mRNA was then fragmented at 94°C for 2 min and copied into first strand complementary DNA (cDNA) using reverse transcriptase and random primers. Second strand cDNA synthesis further generated blunt-ended double-stranded cDNA and incorporated dUTP in place of dTTP to achieve strand specificity by quenching the second strand during amplification. Following A-tailing of DNA fragments and ligation of pre-index anchors, PCR amplification was used to add indexes and primer sequences and to enrich DNA libraries (30 sec at 98°C; [10 sec at 98°C, 30 sec at 60°C, 30 sec at 72°C] x 12 cycles; 5 min at 72°C). Surplus PCR primers were further removed by purification using SPRIselect beads (Beckman-Coulter, Villepinte, France) and the final libraries were checked for quality and quantified using capillary electrophoresis. Libraries were sequenced on an Illumina HiSeq 4000 sequencer as paired-end 100 base reads. Image analysis and base calling were performed using RTA version 2.7.7 and bcl2fastq version 2.20.0.422. Quality check of raw reads was performed using HTseq^64^ and alignment was performed by HISAT2^65^. SAR Tools^66^ was used for differential expression analysis. The functional enrichment analysis of the Differentially Expressed Genes (DEGs) was performed using the web-based tools g:Profiler with default parameters^67^. Heatmap was generated with R package Pheatmap. All Imd/NF-кB pathway related genes which had threshold less than 0.05 (padj < 0.05) were selected and plotted in the figure as Log2 fold change.

#### Immunofluorescence/Confocal imaging and image processing

##### Protocol without antibody staining

Adult fly guts were dissected in ice cold PBS and fixed for 20 min in 4% paraformaldehyde on ice. After 3 washes (5 minutes each) in PBT (1XPBS + 0.1% Triton X-100), the guts were mounted in Vectashield (Vector Laboratories) fluorescent mounting medium, with DAPI. Images of posterior midgut (R4 region) were captured immediately after mounting with a LSM 880 Zeiss confocal microscope, and an individual stack is shown in the figures.

##### Protocol for antibody staining

Following antibodies were used: Primary antibodies: Mouse anti-V5 (1:500, Invitrogen, **R960-25),** Rabbit anti-V5 (1:500, Invitrogen**, MA5-32053),** Rabbit anti-Rab5 (1:500 Abcam, ab31261), Mouse anti-Rab7 (1:500, Abcam, ab50533), Rabbit anti-Lamp1 (1:500, Abcam, ab30687), Chicken anti-GFP (1:2000, Aveslabs, GFP-1020), Mouse anti-Prospero (1:50, DSHB, MR1A). Secondary antibodies: Donkey anti-mouse AF568 (1:500, ThermoFisher Scientific, A10037), Donkey anti-mouse AF647 (1:500, Jackson ImmunoResearch, 715605151), Donkey anti-rabbit AF568 (1:500, ThermoFisher Scientific, A10042), Donkey anti-rabbit AF647 (1:500, ThermoFisher Scientific, A31573). Adult fly guts were dissected in ice cold PBS, fixed for 20 min in 4% paraformaldehyde on ice and washed 3 times in PBT (1XPBS + 0.1% Triton X-100). Tissues were blocked for 2h in PBT + 3% bovine serum albumin (BSA; Sigma-Aldrich) and then incubated overnight with the primary antibodies, in a cold room, on shaker. After 3 washes in PBT, the dissected tissues were incubated 2h with secondary antibodies. The tissues were next rinsed three times in PBT and mounted in Vectashield (Vector Laboratories) fluorescent mounting medium, with DAPI. Images of posterior midgut (R4 region) were captured with a LSM 880 Zeiss confocal microscope, and an individual stack is shown in the all the figures. Some images were false colored for consistency with other images in the manuscript

##### Smurf assay

For Smurf assay 3% (v/v) colorant bleu (E133) was mixed with *E. cc* containing oral infection food. After 6 hpi flies were imaged on Zeiss Stereo Lumar V12 microscope.

#### Data analysis

All images were quantified using Fiji software (available at imagej.nih.gov/ij). The detailed methodology for analysis of various parameters is described as follows.

##### Intensity quantification

Fiji software was used to define the region of interest (ROIs) of 150*150 pixels corresponding to approximately one enterocyte in posterior midgut (R4 region). Random five ROIs were selected using rectangle selection tool keeping nucleus (DAPI) in the center. The mean intensity measurement of PGRP-LE::GFP and PGRP-LE::V5 was then conducted using measure tool (ImageJ function: Analyze > Measure). In all experiments, genotypes were analyzed in parallel. Each experiment was repeated independently at least three times, and one representative quantification is shown. Data were normalized with uninfected and plotted fold change of normalized mean intensity.

##### Clusters number and size quantification

ImageJ macro programs were used to quantify number and size of clusters. Briefly, random five ROIs of 150*150 pixels of single stack were selected using rectangle selection tool keeping nucleus (DAPI) in the center. Next, threshold (ImageJ function: Image > Adjust > Threshold) was set and converted to mask. Watershed (ImageJ function: Process > Binary > Watershed) was used to separate closed clusters by one pixel and then number and size of clusters were quantified with analyze particles tool (ImageJ function: Analyze particles). All the genotypes were analyzed in parallel in all experiments. Each experiment was repeated at least three times, and one representative quantification is shown.

##### Colocalization quantification

ImageJ macro program were used to quantify colocalization. Briefly, random five ROIs of 150*150 pixels of a single stack were selected using rectangle selection tool keeping nucleus (DAPI) in the center. Next, threshold (ImageJ function: Image > Adjust > Threshold) was set and converted to mask. Watershed (ImageJ function: Process > Binary > Watershed) was used to separate closed clusters by one pixel and then number of clusters were quantified with analyze particles tool (ImageJ function: Analyze particles). These quantified clusters were used to quantify colocalization with Rab5 and Rab7 positive endosomes using JACoP (Just Another Colocalization Plugin) plugin in imageJ. Pearson’s R coefficient values were used to plot graphs.

### QUANTIFICATION AND STATISTICAL ANALYSIS

At least three independent repeats were carried out for each experiment (unless otherwise stated in the figure legends). Statistical analyses were conducted comparing independent experiments with use of GraphPad Prism 8 software (GraphPad Software). The total number of animals quantified, p values, and significance levels are indicated in the respective Figure legends.

---

**Figure S1:**
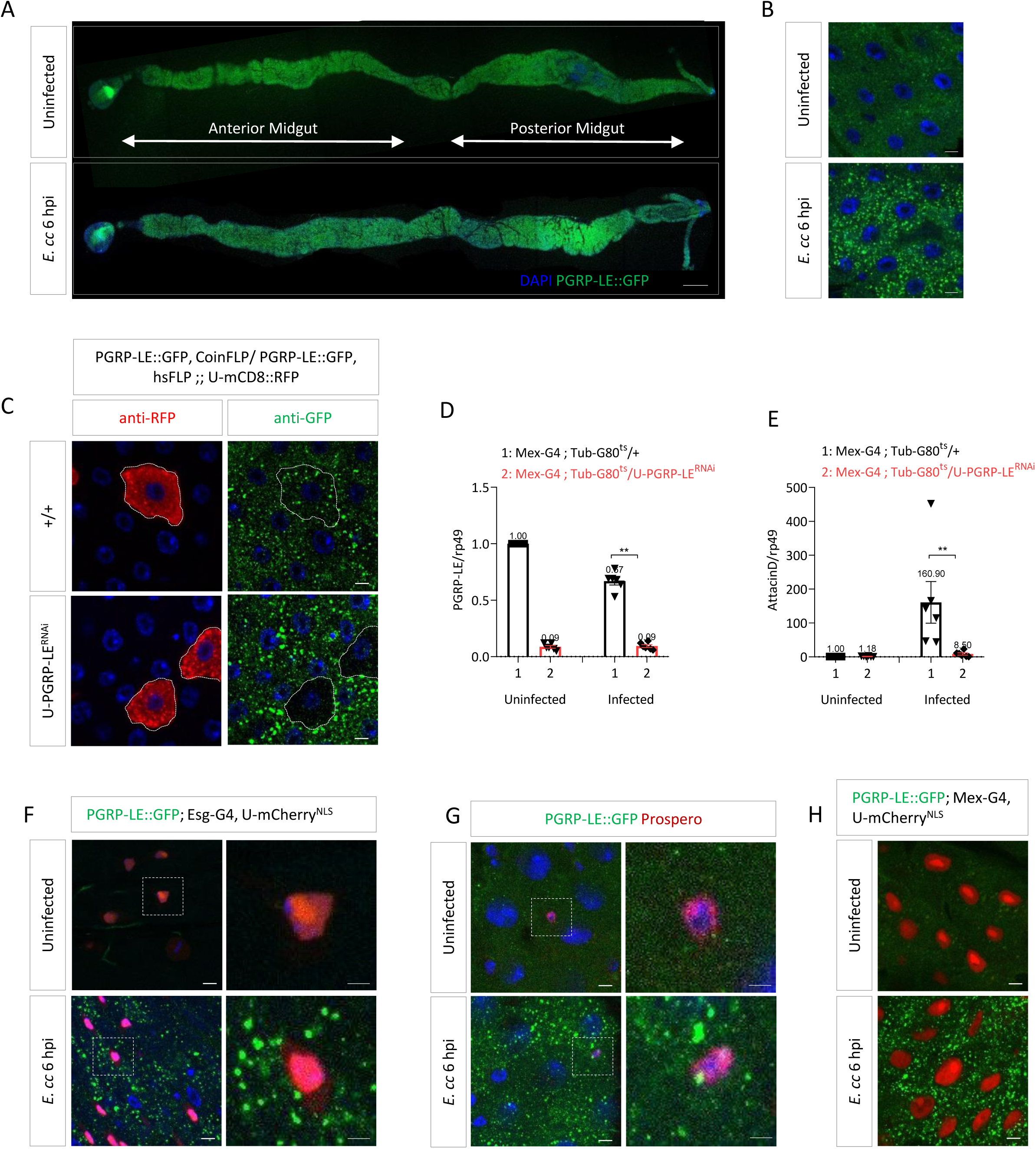
PGRP-LE aggregate characterization, related to Figure 1. **(A)** Adult midguts from uninfected and *E.cc* infected flies. Immunofluorescence showing PGRP-LE::GFP (green) and DNA (blue). Scale bar represents 200 µm. **(B)** R4 region of posterior midguts of uninfected and *E.cc* infected flies showing PGRP-LE::GFP (green) and DNA (blue). Scale bars represent 5 µm. **(C)** Confocal images of posterior midguts from control and PGRP-LE^RNAi^ clones in *E.cc* infected flies showing RFP (clones) in red, PGRP-LE::GFP clusters in green and DNA in blue. Scale bars represent 5 µm. (**D** and **E**) *PGRP-LE* (D) and *AttacinD* (E) mRNA levels in posterior midgut of uninfected and *E.cc* infected flies upon downregulation of PGRP-LE. Data correspond to six independent experiments with ten female flies per genotype per experiment. **p < 0.01; Mann-Whitney test. **(F)** Confocal images of posterior midguts from uninfected and *E.cc* infected flies showing PGRP-LE::GFP (Green) clusters in intestinal stem cells. Nucleus of intestinal stem cells are marked with mCherry. **(G)** Confocal images of posterior midguts from uninfected and *E.cc* infected flies showing PGRP-LE::GFP (Green) clusters in enteroendocrine cells. Nucleus of enteroendocrine cells is marked with prospero. **(H)** Confocal images of posterior midguts from uninfected and *E.cc* infected flies showing PGRP-LE::GFP (Green) clusters in enterocytes. Nucleus of enterocytes are marked with mCherry. Flies were orally infected with *E. cc* for 6 hours. The experiment was repeated three times. Confocal images of one representative experiment are shown from R4 region of posterior midguts. For RT-qPCR results, mRNA levels in uninfected control flies were set to 1, values obtained with other genotypes were expressed as a fold of this value as mean ± SEM.

**Figure S2:**
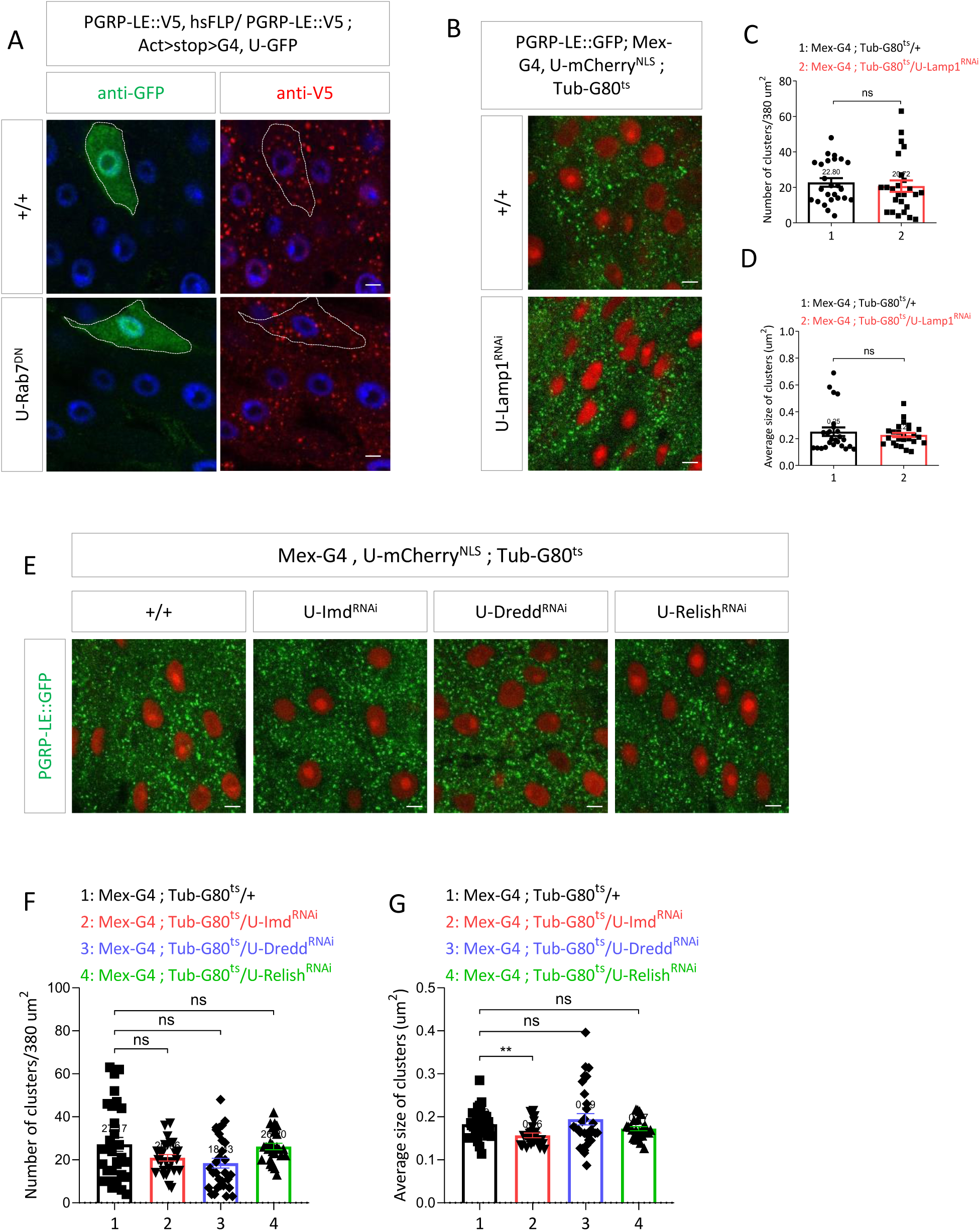
IMD pathway components are not required for *Ecc*-induced PGRP-LE aggregation, related to Figure 4. **(A)** Confocal images of control and Rab7^DN^ clones in posterior midguts of *E.cc* infected flies showing GFP (clones) in green, PGRP-LE::V5 clusters in red and DNA in blue. **(B)** Confocal images of posterior midguts showing PGRP-LE::GFP clusters in control and Lamp1^RNAi^ infected flies. GFP is shown in green and mCherry marks nucleus. (**C** and **D**) Quantification of numbers (C) and average size (D) of PGRP-LE::GFP clusters in control and Lamp1^RNAi^ infected flies with *E. cc* from five posterior midguts. ns (non-significant), p > 0.05; Mann-Whitney test. (**E**) Confocal images of posterior midguts of *E.cc* infected flies showing PGRP-LE::GFP clusters in control, Imd^RNAi^, Dredd^RNAi^ and Relish^RNAi^ flies. GFP is shown in green and mCherry marks nucleus. (**F** and **G**) Quantification of numbers (F) and average size (G) of PGRP-LE::GFP clusters from control, Imd^RNAi^, Dredd^RNAi^ and Relish^RNAi^ infected flies with *E. cc* from five posterior midguts. ns (non-significant), p > 0.05; **p < 0.01; Kruskal-Wallis test. Flies were orally infected with *E. cc* for 6 hours. The experiment was repeated three times. Confocal images of one representative experiment are shown from R4 region of posterior midguts. Scale bars represent 5 µm.

**Figure S3:**
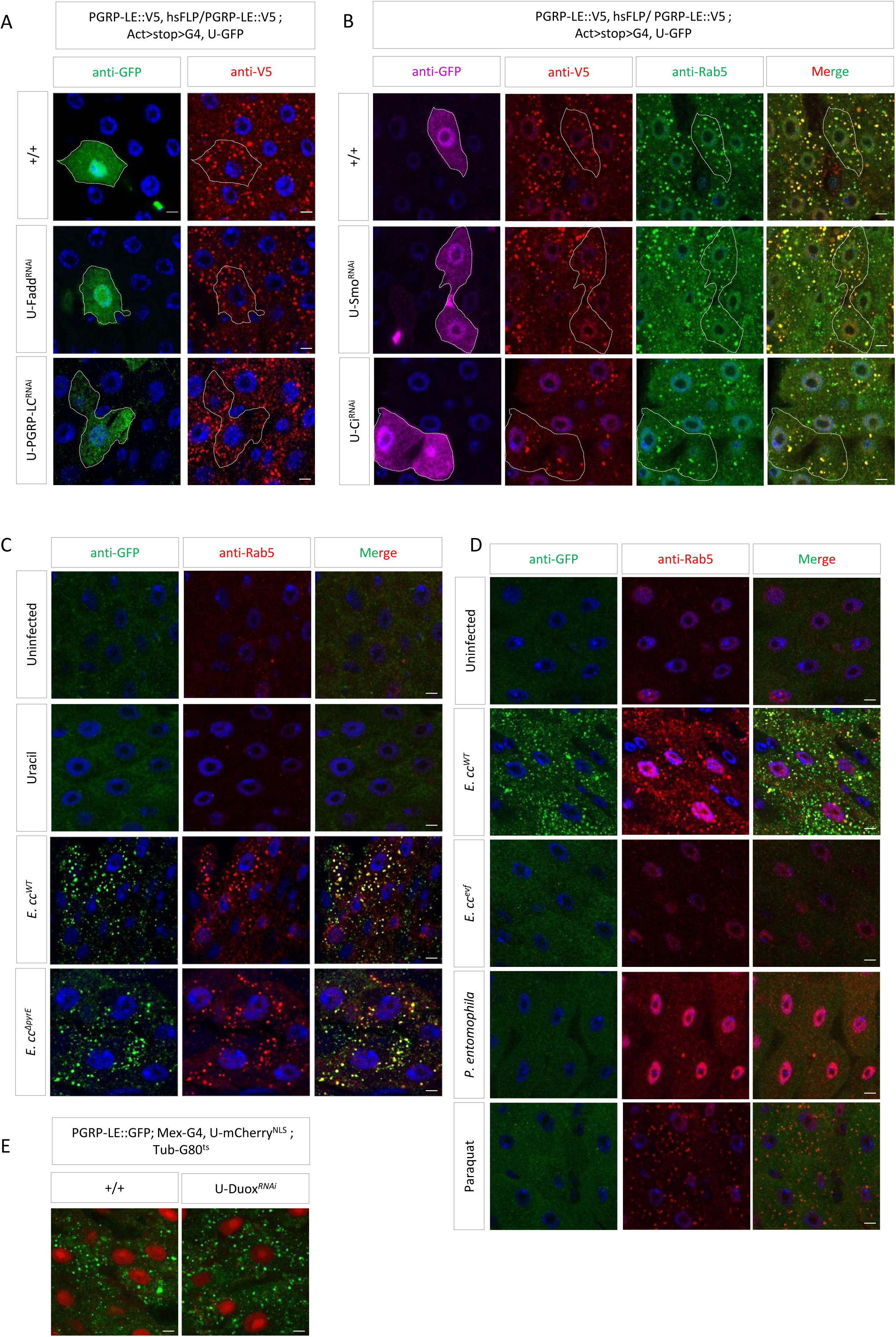
Hh and Duox pathways are not required for *Ecc*-induced PGRP-LE aggregation, related to Figure1. **(A)** Confocal images of control, Fadd^RNAi^ and PGRP-LC^RNAi^ clones in posterior midguts of *E.cc* infected flies. Immunofluorescence showing GFP (clones) in green, PGRP-LE::V5 clusters in red and DNA in blue. **(B)** Confocal images of control, Smo^RNAi^ and Ci^RNAi^ clones of in posterior midguts of *E.cc* infected flies. Immunofluorescence showing GFP (clones) in magenta, PGRP-LE::V5 clusters in red, Rab5 in green and DNA in blue. **(C)** Confocal images of posterior midguts from uninfected flied, flies infected with *E. cc^WT^*, *E. cc*^ΔpyrE^, or flies raised on uracil containing food. **(D)** Confocal images of PGRP-LE::GFP clusters (Green) and Rab5 positive endosomes (Red) from uninfected and infected with *E. cc., E. cc^evf^, P. entomophila* or flies raised on paraquat containing food. **(E)** Confocal images of posterior midguts from of showing PGRP-LE::GFP (Green) clusters in control and Duox^RNAi^ *E.cc*. infected flies. GFP is shown in green and mCherry marks nucleus. Flies were orally infected with the indicated bacteria strains for 6 hours. The experiment was repeated three times. Confocal images of one representative experiment are shown from R4 region of posterior midguts. Scale bars represent 5 µm.

**Figure S4:**
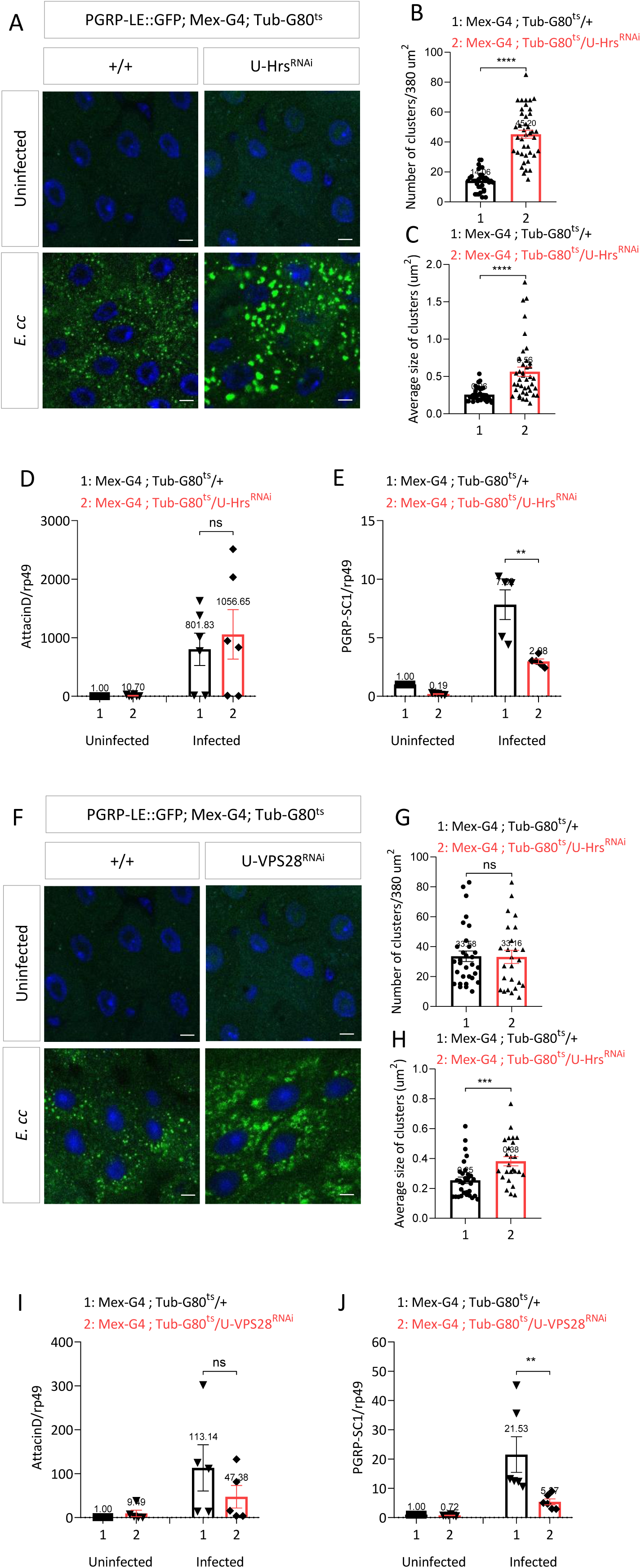
HRS and VPS28 are required for *Ecc*-induced PGRP-LE aggregation, related to Figure 6. **(A)** Confocal images of posterior midguts showing PGRP-LE::GFP clusters in control and Hrs^RNAi^ infected flies. GFP is shown in green and DNA in blue. (**B** and **C**) Quantification of numbers (B) and average size (C) of PGRP-LE::GFP clusters in control and Hrs^RNAi^ infected flies with *E. cc* from seven posterior midguts. **p < 0.01; ****p < 0.0001; Mann-Whitney test. (**D** and **E**) *AttacinD* (D) and *PGRP-SC1* (E) mRNA levels in posterior midgut of uninfected and *E.cc* infected flies upon downregulation of Hrs. Data correspond to five independent experiments with ten female flies per genotype per experiment. ns (non-significant), p > 0.05; **p < 0.01; Mann-Whitney test. **(F)** Confocal images of posterior midguts showing PGRP-LE::GFP clusters in control and VPS28^RNAi^ infected flies. GFP is shown in green and DNA in blue. (**G** and **H**) Quantification of numbers (G) and average size (H) of PGRP-LE::GFP clusters in control and VPS28^RNAi^ infected flies with *E. cc* from five posterior midguts. ns (non-significant), p > 0.05; ***p < 0.001; Mann-Whitney test. (**I** and **J**) *AttacinD* (I) and *PGRP-SC1* (J) mRNA levels in posterior midgut of uninfected and *E.cc* infected flies upon downregulation of VPS28. Data correspond to five independent experiments with ten female flies per genotype per experiment. ns (non-significant), p > 0.05; **p < 0.01; Mann-Whitney test. Flies were orally infected with the indicated bacteria strains for 6 hours. The experiment was repeated three times. Confocal images of one representative experiment are shown from R4 region of posterior midguts. Scale bars represent 5 µm. For RT-qPCR results, mRNA levels in uninfected control flies were set to 1, values obtained with other genotype were expressed as a fold of this value. RT-qPCR results are shown as mean ± SEM.

## References

1. Round, J.L., and Mazmanian, S.K. (2009). The gut microbiota shapes intestinal immune responses during health and disease. Nature Reviews Immunology 2009 9:5 9, 313–323. 10.1038/NRI2515.

2. Kawai, T., and Akira, S. (2009). The roles of TLRs, RLRs and NLRs in pathogen recognition. Int Immunol 21, 317–337. 10.1093/INTIMM/DXP017.

3. Takeuchi, O., and Akira, S. (2010). Pattern Recognition Receptors and Inflammation. Cell 140, 805–820. 10.1016/J.CELL.2010.01.022.

4. Bastos, P.A.D., Wheeler, R., and Boneca, I.G. (2021). Uptake, recognition and responses to peptidoglycan in the mammalian host. FEMS Microbiol Rev 45, 1–25. 10.1093/FEMSRE/FUAA044.

5. Tosoni, G., Conti, M., and Diaz Heijtz, R. (2019). Bacterial peptidoglycans as novel signaling molecules from microbiota to brain. Curr Opin Pharmacol 48, 107–113. 10.1016/J.COPH.2019.08.003.

6. Wolf, A.J. (2023). Peptidoglycan-induced modulation of metabolic and inflammatory responses. Immunometabolism (Cobham (Surrey, England) 5, e00024. 10.1097/IN9.0000000000000024.

7. Irazoki, O., Hernandez, S.B., and Cava, F. (2019). Peptidoglycan muropeptides: Release, perception, and functions as signaling molecules. Front Microbiol 10, 500. 10.3389/FMICB.2019.00500/BIBTEX.

8. Girardin, S.E., Boneca, I.G., Carneiro, L.A.M., Antignac, A., Jéhanno, M., Viala, J., Tedin, K., Taha, M.K., Labigne, A., Zähringer, U., et al. (2003). Nod1 detects a unique muropeptide from gram-negative bacterial peptidoglycan. Science 300, 1584–1587. 10.1126/SCIENCE.1084677.

9. Schlöffel, M.A., Käsbauer, C., and Gust, A.A. (2019). Interplay of plant glycan hydrolases and LysM proteins in plant—Bacteria interactions. International Journal of Medical Microbiology 309, 252–257. 10.1016/J.IJMM.2019.04.004.

10. Mesnage, S., Dellarole, M., Baxter, N.J., Rouget, J.B., Dimitrov, J.D., Wang, N., Fujimoto, Y., Hounslow, A.M., Lacroix-Desmazes, S., Fukase, K., et al. (2014). Molecular basis for bacterial peptidoglycan recognition by LysM domains. Nat Commun 5. 10.1038/NCOMMS5269.

11. Saur, I.M.L., Panstruga, R., and Schulze-Lefert, P. (2020). NOD-like receptor-mediated plant immunity: from structure to cell death. Nature Reviews Immunology 2020 21:5 21, 305–318. 10.1038/S41577-020-00473-Z.

12. Royet, J., Gupta, D., and Dziarski, R. (2011). Peptidoglycan recognition proteins: modulators of the microbiome and inflammation. Nat Rev Immunol 11, 837–851. 10.1038/NRI3089.

13. Grenier, T., and Leulier, F. (2020). How commensal microbes shape the physiology of Drosophila melanogaster. Curr Opin Insect Sci 41, 92–99. 10.1016/J.COIS.2020.08.002.

14. Buchon, N., Silverman, N., and Cherry, S. (2014). Immunity in Drosophila melanogaster — from microbial recognition to whole-organism physiology. Nature Reviews Immunology 2014 14:12 14, 796–810. 10.1038/NRI3763.

15. Broderick, N.A., and Lemaitre, B. (2012). Gut-associated microbes of Drosophila melanogaster. Gut Microbes 3. 10.4161/GMIC.19896.

16. Lestradet, M., Lee, K.Z., and Ferrandon, D. (2014). Drosophila as a model for intestinal infections. Methods in Molecular Biology 1197, 11–40. 10.1007/978-1-4939-1261-2_2/COVER.

17. Schneider, J., and Imler, J.L. (2021). Sensing and signalling viral infection in drosophila. Dev Comp Immunol 117, 103985. 10.1016/J.DCI.2020.103985.

18. Charroux, B., Capo, F., Kurz, C.L., Peslier, S., Chaduli, D., Viallat-lieutaud, A., and Royet, J. (2018). Cytosolic and Secreted Peptidoglycan-Degrading Enzymes in Drosophila Respectively Control Local and Systemic Immune Responses to Microbiota. Cell Host Microbe 23, 215–228.e4. 10.1016/j.chom.2017.12.007.

19. Capo, F., Charroux, B., and Royet, J. (2016). Bacteria sensing mechanisms in Drosophila gut: Local and systemic consequences. Dev Comp Immunol 64, 11–21. 10.1016/J.DCI.2016.01.001.

20. Bosco-Drayon, V., Poidevin, M., Boneca, I.G., Narbonne-Reveau, K., Royet, J., and Charroux, B. (2012). Peptidoglycan sensing by the receptor PGRP-LE in the Drosophila gut induces immune responses to infectious bacteria and tolerance to microbiota. Cell Host Microbe 12, 153–165. 10.1016/j.chom.2012.06.002.

21. Leulier, F., Parquet, C., Pili-Floury, S., Ryu, J.H., Caroff, M., Lee, W.J., Mengin-Lecreulx, D., and Lemaitre, B. (2003). The Drosophila immune system detects bacteria through specific peptidoglycan recognition. Nature Immunology 2003 4:5 4, 478–484. 10.1038/NI922.

22. Gendrin, M., Welchman, D.P., Poidevin, M., Hervé, M., and Lemaitre, B. (2009). Long-Range Activation of Systemic Immunity through Peptidoglycan Diffusion in Drosophila. PLoS Pathog 5, e1000694. 10.1371/JOURNAL.PPAT.1000694.

23. Kleino, A., Ramia, N.F., Bozkurt, G., Shen, Y., Nailwal, H., Huang, J., Napetschnig, J., Gangloff, M., Chan, F.K.M., Wu, H., et al. (2017). Peptidoglycan-Sensing Receptors Trigger the Formation of Functional Amyloids of the Adaptor Protein Imd to Initiate Drosophila NF-κB Signaling. Immunity 47, 635–647.e6. 10.1016/j.immuni.2017.09.011.

24. Neyen, C., Runchel, C., Schüpfer, F., Meier, P., and Lemaitre, B. (2016). The regulatory isoform rPGRP-LC induces immune resolution via endosomal degradation of receptors. Nature Immunology 2016 17:10 17, 1150–1158. 10.1038/NI.3536.

25. Maillet, F., Bischoff, V., Vignal, C., Hoffmann, J., and Royet, J. (2008). The Drosophila Peptidoglycan Recognition Protein PGRP-LF Blocks PGRP-LC and IMD/JNK Pathway Activation. Cell Host Microbe 3, 293–303. 10.1016/j.chom.2008.04.002.

26. Gottar, M., Gobert, V., Michel, T., Belvin, M., Duyk, G., Hoffmann, J.A., Ferrandon, D., and Royet, J. (2002). The Drosophila immune response against Gram-negative bacteria is mediated by a peptidoglycan recognition protein. Nature 2002 416:6881 416, 640–644. 10.1038/NATURE734.

27. Choe, K.M., Werner, T., Stöven, S., Hultmark, D., and Anderson, K. V. (2002). Requirement for a peptidoglycan recognition protein (PGRP) in relish activation and antibacterial immune responses in Drosophila. Science (1979) 296, 359–362. 10.1126/SCIENCE.1070216/SUPPL_FILE/1070216S1_THUMB.GIF.

28. Kurata, S. (2010). Extracellular and intracellular pathogen recognition by Drosophila PGRP-LE and PGRP-LC. Int Immunol 22, 143–148. 10.1093/INTIMM/DXP128.

29. Yano, T., Mita, S., Ohmori, H., Oshima, Y., Fujimoto, Y., Ueda, R., Takada, H., Goldman, W.E., Fukase, K., Silverman, N., et al. (2008). Autophagic control of listeria through intracellular innate immune recognition in drosophila. Nature Immunology 2008 9:8 9, 908–916. 10.1038/NI.1634.

30. Kaneko, T., Yano, T., Aggarwal, K., Lim, J.H., Ueda, K., Oshima, Y., Peach, C., Erturk-Hasdemir, D., Goldman, W.E., Oh, B.H., et al. (2006). PGRP-LC and PGRP-LE have essential yet distinct functions in the drosophila immune response to monomeric DAP-type peptidoglycan. Nature Immunology 2006 7:7 7, 715–723. 10.1038/NI1356.

31. Takehana, A., Yano, T., Mita, S., Kotani, A., Oshima, Y., and Kurata, S. (2004). Peptidoglycan recognition protein (PGRP)-LE and PGRP-LC act synergistically in Drosophila immunity. EMBO J 23, 4690–4700. 10.1038/SJ.EMBOJ.7600466.

32. Chevée, V., Sachar, U., Yadav, S., Heryanto, C., and Eleftherianos, I. (2019). The peptidoglycan recognition protein PGRP-LE regulates the Drosophila immune response against the pathogen Photorhabdus. Microb Pathog 136, 103664. 10.1016/J.MICPATH.2019.103664.

33. Charroux, B., and Royet, J. (2022). Gut-derived peptidoglycan remotely inhibits bacteria dependent activation of SREBP by Drosophila adipocytes. PLoS Genet 18, e1010098. 10.1371/JOURNAL.PGEN.1010098.

34. Masuzzo, A., Montanari, M., Kurz, L., and Royet, J. (2020). How Bacteria Impact Host Nervous System and Behaviors: Lessons from Flies and Worms. Trends Neurosci 43, 998– 1010. 10.1016/j.tins.2020.09.007.

35. Masuzzo, A., Manière, G., Viallat-Lieutaud, A., Avazeri, É., Zugasti, O., Grosjean, Y., Kurz, C.L., and Royet, J. (2019). Peptidoglycan-dependent NF-kB activation in a small subset of brain octopaminergic neurons controls female oviposition. Elife 8. 10.7554/ELIFE.50559.

36. Kurz, C.L., Charroux, B., Chaduli, D., Viallat-Lieutaud, A., and Royet, J. (2017). Peptidoglycan sensing by octopaminergic neurons modulates Drosophila oviposition. Elife 6. 10.7554/ELIFE.21937.

37. Kleino, A., and Silverman, N. (2014). The Drosophila IMD pathway in the activation of the humoral immune response. Dev Comp Immunol 42, 25–35. 10.1016/J.DCI.2013.05.014.

38. Tzou, P., Ohresser, S., Ferrandon, D., Capovilla, M., Reichhart, J.M., Lemaitre, B., Hoffmann, J.A., and Imler, J.L. (2000). Tissue-specific inducible expression of antimicrobial peptide genes in Drosophila surface epithelia. Immunity 13, 737–748. 10.1016/S1074-7613(00)00072-8.

39. Irving, A.T., Mimuro, H., Kufer, T.A., Lo, C., Wheeler, R., Turner, L.J., Thomas, B.J., Malosse, C., Gantier, M.P., Casillas, L.N., et al. (2014). The immune receptor NOD1 and kinase RIP2 interact with bacterial peptidoglycan on early endosomes to promote autophagy and inflammatory signaling. Cell Host Microbe 15, 623–635. 10.1016/j.chom.2014.04.001.

40. Nakamura, N., Lill, J.R., Phung, Q., Jiang, Z., Bakalarski, C., De Mazière, A., Klumperman, J., Schlatter, M., Delamarre, L., and Mellman, I. (2014). Endosomes are specialized platforms for bacterial sensing and NOD2 signalling. Nature 2014 509:7499 509, 240–244. 10.1038/NATURE13133.

41. Basset, A., Khush, R.S., Braun, A., Gardan, L., Boccard, F., Hoffmann, J.A., and Lemaitre, B. (2000). The phytopathogenic bacteria Erwinia carotovora infects Drosophila and activates an immune response. Proc Natl Acad Sci U S A 97, 3376–3381. 10.1073/PNAS.97.7.3376/ASSET/ED735D10-9886-4947-AAA7-3C2AF0FB6C19/ASSETS/GRAPHIC/PQ0703575005.JPEG.

42. Storelli, G., Defaye, A., Erkosar, B., Hols, P., Royet, J., and Leulier, F. (2011). Lactobacillus plantarum promotes drosophila systemic growth by modulating hormonal signals through TOR-dependent nutrient sensing. Cell Metab 14, 403–414. 10.1016/j.cmet.2011.07.012.

43. Shin, S.C., Kim, S.H., You, H., Kim, B., Kim, A.C., Lee, K.A., Yoon, J.H., Ryu, J.H., and Lee, W.J. (2011). Drosophila microbiome modulates host developmental and metabolic homeostasis via insulin signaling. Science (1979) 334, 670–674. 10.1126/SCIENCE.1212782/SUPPL_FILE/1212782.SHIN.SOM.PDF.

44. Liehl, P., Blight, M., Vodovar, N., Boccard, F., and Lemaitre, B. (2006). Prevalence of Local Immune Response against Oral Infection in a Drosophila/Pseudomonas Infection Model. PLoS Pathog 2, e56. 10.1371/JOURNAL.PPAT.0020056.

45. Muniz, C.A., Jaillard, D., Lemaitre, B., and Boccard, F. (2007). Erwinia carotovora Evf antagonizes the elimination of bacteria in the gut of Drosophila larvae. Cell Microbiol 9, 106–119. 10.1111/J.1462-5822.2006.00771.X.

46. Lim, J.H., Kim, M.S., Kim, H.E., Yano, T., Oshima, Y., Aggarwal, K., Goldman, W.E., Silverman, N., Kurata, S., and Oh, B.H. (2006). Structural basis for preferential recognition of diaminopimelic acid-type peptidoglycan by a subset of peptidoglycan recognition proteins. Journal of Biological Chemistry 281, 8286–8295. 10.1074/jbc.M513030200.

47. Bonham, K.S., and Kagan, J.C. (2014). Endosomes as platforms for NOD-like receptor signaling. Cell Host Microbe 15, 523–525. 10.1016/j.chom.2014.05.001.

48. Clague, M.J., and Urbé, S. (2001). The interface of receptor trafficking and signalling. J Cell Sci 114, 3075–3081. 10.1242/JCS.114.17.3075.

49. Jacomin, A.C., Fauvarque, M.O., and Taillebourg, E. (2016). A functional endosomal pathway is necessary for lysosome biogenesis in Drosophila. BMC Cell Biol 17, 1–9. 10.1186/S12860-016-0115-7/FIGURES/5.

50. Lee, K.A., Kim, B., Bhin, J., Kim, D.H., You, H., Kim, E.K., Kim, S.H., Ryu, J.H., Hwang, D., and Lee, W.J. (2015). Bacterial uracil modulates drosophila DUOX-dependent Gut immunity via hedgehog-induced signaling endosomes. Cell Host Microbe 17, 191–204. 10.1016/j.chom.2014.12.012.

51. Maitra, U., Scaglione, M.N., Chtarbanova, S., and O’Donnell, J.M. (2019). Innate immune responses to paraquat exposure in a Drosophila model of Parkinson’s disease. Sci Rep 9. 10.1038/S41598-019-48977-6.

52. Zaidman-Rémy, A., Hervé, M., Poidevin, M., Pili-Floury, S., Kim, M.S., Blanot, D., Oh, B.H., Ueda, R., Mengin-Lecreulx, D., and Lemaitre, B. (2006). The Drosophila Amidase PGRP-LB Modulates the Immune Response to Bacterial Infection. Immunity 24, 463–473. 10.1016/j.immuni.2006.02.012.

53. Rera, M., Clark, R.I., and Walker, D.W. (2012). Intestinal barrier dysfunction links metabolic and inflammatory markers of aging to death in Drosophila. Proc Natl Acad Sci U S A 109, 21528–21533. 10.1073/PNAS.1215849110/SUPPL_FILE/SM01.AVI.

54. Onuma, T., Yamauchi, T., Kosakamoto, H., Kadoguchi, H., Kuraishi, T., Murakami, T., Mori, H., Miura, M., and Obata, F. (2023). Recognition of commensal bacterial peptidoglycans defines Drosophila gut homeostasis and lifespan. PLoS Genet 19, e1010709. 10.1371/JOURNAL.PGEN.1010709.

55. Huang, H.R., Chen, Z.J., Kunes, S., Chang, G.D., and Maniatis, T. (2010). Endocytic pathway is required for Drosophila Toll innate immune signaling. Proc Natl Acad Sci U S A 107, 8322– 8327. 10.1073/PNAS.1004031107/SUPPL_FILE/PNAS.201004031SI.PDF.

56. Dixit, E., Boulant, S., Zhang, Y., Lee, A.S.Y., Odendall, C., Shum, B., Hacohen, N., Chen, Z.J., Whelan, S.P., Fransen, M., et al. (2010). Peroxisomes Are Signaling Platforms for Antiviral Innate Immunity. Cell 141, 668–681. 10.1016/j.cell.2010.04.018.

57. Banoth, B., and Cassel, S.L. (2018). Mitochondria in innate immune signaling. Translational Research 202, 52–68. 10.1016/J.TRSL.2018.07.014.

58. Sorvina, A., Shandala, T., and Brooks, D.A. (2016). Drosophila Pkaap regulates Rab4/Rab11-dependent traffic and Rab11 exocytosis of innate immune cargo. Biol Open 5, 678–688. 10.1242/BIO.016642/-/DC1.

59. Zhang, P., Holowatyj, A.N., Roy, T., Pronovost, S.M., Marchetti, M., Liu, H., Ulrich, C.M., and Edgar, B.A. (2019). An SH3PX1-Dependent Endocytosis-Autophagy Network Restrains Intestinal Stem Cell Proliferation by Counteracting EGFR-ERK Signaling. Dev Cell 49, 574–589.e5. 10.1016/j.devcel.2019.03.029.

60. Nagy, P., Kovács, L., Sándor, G.O., and Juhász, G. (2016). Stem-cell-specific endocytic degradation defects lead to intestinal dysplasia in Drosophila. DMM Disease Models and Mechanisms 9, 501–512. 10.1242/DMM.023416/257176/AM/STEM-CELL-SPECIFIC-ENDOCYTIC-DEGRADATION-DEFECTS.

61. Nassari, S., Lacarrière-Keïta, C., Lévesque, D., Boisvert, F.M., and Jean, S. (2022). Rab21 in enterocytes participates in intestinal epithelium maintenance. Mol Biol Cell 33. 10.1091/MBC.E21-03-0139/ASSET/IMAGES/LARGE/MBC-33-AR32-G009.JPEG.

62. Tang, R., Huang, W., Guan, J., Liu, Q., Beerntsen, B.T., and Ling, E. (2021). Drosophila H2Av negatively regulates the activity of the IMD pathway via facilitating Relish SUMOylation. PLoS Genet 17, e1009718. 10.1371/JOURNAL.PGEN.1009718.

63. Kane, N.S., Vora, M., Varre, K.J., and Padgett, R.W. (2017). Efficient screening of CRISPR/Cas9-induced events in Drosophila using a Co-CRISPR strategy. G3: Genes, Genomes, Genetics 7, 87–93. 10.1534/G3.116.036723/-/DC1.

64. Anders, S., Pyl, P.T., and Huber, W. (2015). HTSeq—a Python framework to work with high-throughput sequencing data. Bioinformatics 31, 166–169. 10.1093/BIOINFORMATICS/BTU638.

65. Kim, D., Paggi, J.M., Park, C., Bennett, C., and Salzberg, S.L. (2019). Graph-based genome alignment and genotyping with HISAT2 and HISAT-genotype. Nature Biotechnology 2019 37:8 37, 907–915. 10.1038/S41587-019-0201-4.

66. Varet, H., Brillet-Guéguen, L., Coppée, J.Y., and Dillies, M.A. (2016). SARTools: A DESeq2- and EdgeR-Based R Pipeline for Comprehensive Differential Analysis of RNA-Seq Data. PLoS One 11, e0157022. 10.1371/JOURNAL.PONE.0157022.

67. Raudvere, U., Kolberg, L., Kuzmin, I., Arak, T., Adler, P., Peterson, H., and Vilo, J. (2019). g:Profiler: a web server for functional enrichment analysis and conversions of gene lists (2019 update). Nucleic Acids Res 47, W191–W198. 10.1093/NAR/GKZ369.

